# Single-cell characterization of skin response to a bite by West Nile virus-infected mosquito reveals fibroblast-mediated barrier to transmission

**DOI:** 10.64898/2026.01.08.698384

**Authors:** Hacène Medkour, Federico Bocci, Norman Schneider, Idalba Serrato-Pomar, Félix Rey-Cadilhac, Yingzi Liu, Elliott F. Miot, Axel A. Almet, Marine Louarn, Nicolas Aubourg, Wilfried Saron, Ghizlane Maarifi, Dorothée Missé, Qing Nie, Duncan R. Smith, Sébastien Nisole, Ashley St John, Charles Dutertre, Maksim V. Plikus, Julien Pompon

## Abstract

Cutaneous events at the mosquito bite site that determine orthoflavivirus transmission efficiency remain largely uncharacterized. Here, we report single-cell RNA-sequencing of skin from immunocompetent mice bitten by West Nile virus-infected mosquitoes, capturing early response at a critical transmission bottleneck. Fibroblasts were the dominant cells exposed to infectious saliva. While neutrophils and *Folr2*^+^ macrophages diminished, monocyte-derived and antigen-presenting macrophages, and lymphocytes became more abundant. Cell-cell communication analysis revealed the central roles of fibroblasts and myeloid cells in induced immune-related signaling. Transcriptional profiling defined cell type-specific responses to infectious bite integrating immune modulation, skin repair and metabolic remodeling. Using *in vivo* gene silencing, we demonstrated that fibroblast-expressed LRRC15 (leucine rich repeat-containing 15) functions as a cutaneous restriction factor, limiting viral replication in skin, viral dissemination to draining lymph nodes, and disease severity. Collectively, our analyses provide cellular and molecular understanding of bite-initiated arboviral transmission, establishing skin-resident fibroblasts as frontline defender cells.

**HIGHLIGHTS:**

- Single-cell profiling captures early cutaneous response to West Nile virus-infected mosquito bites with cell type resolution.
- Skin-resident fibroblasts are the dominant frontline cells exposed to infectious mosquito saliva.
- Infectious bite reconfigures cutaneous immune cells, and activates multi-directional signaling between immune and structural skin cell types.
- Bite-induced fibroblast-expressed LRRC15 restricts viral transmission and attenuates disease severity.

## INTRODUCTION

Mosquito-borne orthoflaviviruses—including West Nile virus (WNV), dengue virus (DENV) and Zika virus (ZIKV)—cause recurrent outbreaks worldwide, infecting an estimated half a billion people each year and imposing considerable health and economic burden^1,2^. Existing vaccines remain limited and raise safety concerns, complicating both their design and clinical evaluation^3–6^. No antiviral drugs are licensed to-date and therapeutic approaches for acute infection are constrained by an extremely narrow treatment window^7,8^. These challenges underscore the urgent need for novel intervention strategies for mosquito-transmitted infectious diseases.

Bite-initiated skin infection is a critical vulnerability in viral transmission, representing a potential therapeutic target^9,10^. During probing, virus-laden mosquito saliva is injected into the skin, initiating viral transmission through a coordinated molecular and cellular response^11–13^. Saliva and bite-triggered mechanical injury induce local tissue inflammation and increase vascular permeability, which together promote recruitment of neutrophils. These neutrophils release chemoattractants that then drive tissue infiltration by monocytes and the activation or repositioning of resident dendritic cells (DC) and macrophages^14–21^. These infiltrating myeloid cells—themselves being highly susceptible to orthoflavivirus infection—fulfill their defensive immune function as antigen-presenting cells while concurrently carrying viruses to draining lymph nodes (LNs), and as such, enable systemic dissemination and disease transmission^16,17,21–25^. Bite-induced skin edema further facilitates initial virus replication by restricting virus dispersion and maintaining saliva-delivered inoculum^11,17,19^. This sequence of events unfolds with remarkable speed: transcriptional up-regulation of neutrophil-attracting chemokines and neutrophil influx occurs within 4 hours post-bite, myeloid cell infiltration by 6^th^ hour^16,19^ and viral spread to LNs within 12-24 hours^11,25–27^. Consistently, excision of the bitten skin site at 4 hours or therapeutic activation of antiviral defenses before 5^th^ hour post-bite abolishes transmission^21,28^.

Structural cells of the skin also contribute to these early events. Major structural skin cells—keratinocytes in the epidermis and ectodermal appendages, as well as fibroblasts in the dermis^29^—are likely among the first cells to encounter infectious mosquito saliva and their permissiveness to multiple orthoflaviviruses may influence disease transmission^20,30–33^. Upon infection, structural cells activate innate immune responses^32,34–37^, including interferon (IFN) pathways, which limit viral replication^33^. Moreover, infected fibroblasts promote DCs’ viral resistance and T cell recruitment^31,38^, highlighting the importance of cell-cell communication in the cutaneous response to infectious bites.

Despite these insights, prevailing mechanistic models of bite-initiated viral transmission rely largely on indirect and approximate experimental systems—such as *ex vivo* virus incubation of skin explants or dissociated skin cells, artificial needle-based injection of viruses either without or with saliva or with mosquito salivary gland extracts into immunodeficient mice, uninfected bites followed by viral needle-based inoculation or uninfected bite alone^25,39–41^. However, the well-documented impact of mosquito bite on viral transmission kinetics, tissue tropism, cutaneous gene expression and disease severity^17,42,43^ calls for the use of an *in vivo* transmission model to study the early events of mosquito-borne viral transmission. Here, for the first time we applied single-cell RNA-sequencing (scRNA-seq) to skin of an immunocompetent mouse model exposed to infectious mosquito bites to characterize the early molecular and cellular responses to viral transmission. Our study identified the specific cell types that first encounter infectious saliva, delineated the rapid reorganization of the cellular landscape of the skin, mapped induced cell-cell communications and defined cell subtype-specific transcriptomic responses. Finally, we revealed a fibroblast-mediated bite-induced antiviral program that restricts viral transmission.

## RESULTS

### Single-cell characterization of mouse skin bitten by infectious mosquitoes

To conduct informative scRNA-seq evaluation of infectious mosquito-bitten mouse skin, we first characterized the *in vivo* bite-initiated transmission dynamics in wild-type (WT) mice. Multiple mosquitoes infected with WNV by intrathoracic inoculation—to standardize the inoculum—were allowed to bite immunocompetent WT mice (Fig. S1A). All mosquitoes had infected salivary glands (Fig. S1B,C) and their bites induced viremia (peaking at 4 days post-bite), clinical signs of infection, prominent weight loss and mortality in experimental mice (Fig. S1D-G; Dataset S1A,B), thereby validating viral transmission. To define an optimal time point for the scRNA-seq assay, we next characterized viral infection kinetics in the bitten skin. We developed a method for precise identification of the bite site: individually caged mosquitoes (as described in^13^) (Fig. S2A,B) were placed on shaved mouse abdomen and access to the skin was further directed by applying tape perforated with four 2 mm-diameter openings (Fig. S2C,D). Visually identified bite sites were marked and collected from 10 minutes to 24 hours post-bite (Fig. S2E). Because the mosquito proboscis penetrates < 2 mm in any direction into mouse skin^13^, we collected 2-mm-diameter skin biopsies and used them to quantify WNV genomic RNA (gRNA) (Fig. 1A). At 10 minutes post-bite, before virus amplification^44,45^, ∼10^4^ copies of gRNA were expectorated into the skin. This gRNA level remained stable until 3 hours post-bite (hpb), then declined at 6-to-9 hpb—matching the time of viral genome replication onset^44,45^—before increasing to ∼9.6 x 10^4^ copies at 24 hpb, indicating virus multiplication. To verify dissemination, WNV gRNA was quantified in draining LNs (Fig. 1B). Viral gRNA was detected as early as 10 minutes post-bite (geometric mean, 345 gRNA copies per LN), remained stable until 9 hpb, and then sharply increased to ∼4.1 x 10^4^ copies at 24 hpb, validating systemic infection. Based on these kinetics, we selected 6 hpb for cutaneous scRNA-seq analysis, a time point corresponding to the bottleneck phase characterized by transient reduction in cutaneous viral load and the absence of detectable viral expansion in draining LNs.

**Figure 1.**
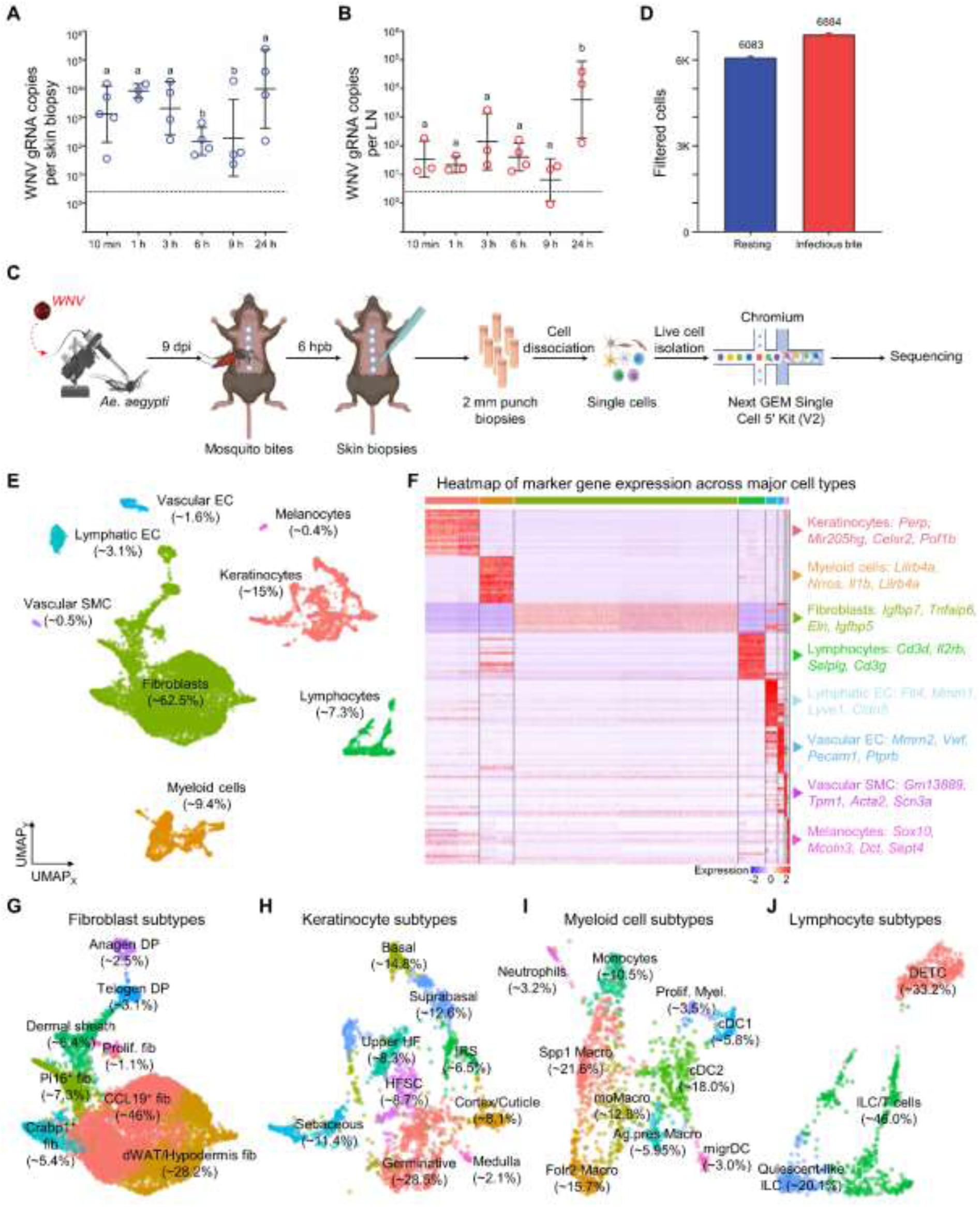
scRNA-seq analysis on mouse skin at the onset of bite-initiated infection. **(A,B)** Early WNV replication kinetics in the skin (A) and inguinal LNs of mice (B) at 10 min, 1, 3, 6, 9 and 24 hours (h) post-bite. Bars show geometric means ± 95% C.I. Point represents one skin biopsy (A) and the two inguinal LNs from the corresponding mouse **(B)**. N mice, 3-4 per time point. Different letters indicate statistical differences as determined by post-hoc Fisher LSD’s test. **(C)** Experimental workflow for the scRNA-seq assay on bitten mouse skin. **(D)** Numbers of high-quality single cells retained after quality control (QC) on scRNA-seq data. **(E)** UMAP dimensional reduction of sequenced skin cells revealing eight cell types. Proportion of each cell cluster is provided in brackets. EC, endothelial cells; SMC, smooth muscle cells. **(F)** Heatmap of the top 20 enriched genes per cell cluster. Four marker genes for each cluster are shown on the right. **(G-J)** UMAP subclustering of fibroblasts (G), keratinocytes and ectodermal appendage cells (H), myeloid cells (I), and lymphocytes (J). Relative cell proportions are indicated in brackets. Fib, fibroblasts; Ker, keratinocytes; DP, dermal papilla cells; dWAT, dermal white adipose tissue cells; HF, hair follicle cells; IRS, inner root sheath cells; HFSC, hair follicle stem cells; DETC, dendritic epidermal T cells; ILC, innate lymphoid T cells. See also **Figures S1-S6 and Datasets S1A-B, S2, S3.**

To perform scRNA-seq on skin at 6 hpb (Fig. 1C), we collected 2-mm biopsies centered on the bite sites to enrich for cells directly exposed to infectious saliva and processed tissues for single-cell isolation and capture. To minimize inter-sample variability and to reduce animal use, each mouse received between 3 and 5 bites, yielding multiple skin biopsies per individual. Multiple bites did not alter local cutaneous viral load at 24 hpb (Fig. S1H) but increased the viral inoculum reaching draining LNs (Fig. S1I). We optimized skin cell dissociation using enzymatic digestion combined with needle-assisted mechanical disruption to improve recovery of viable cells, including dermal DCs (Fig. S3A-C). Skin from non-bitten WT mice (hereafter referred to as resting skin) was identically processed and served as a control. After sequencing, genome mapping, quality controls and bioinformatic sample integration (Fig. S4A-E), we obtained > 6,000 high quality sequenced cells per experimental condition (Fig. 1D).

Unsupervised clustering identified eight cell types (Fig. 1E), each defined by a distinct transcriptional signature (Fig. 1F; Dataset S2), including established cell type-specific marker genes (Fig. S5A-H). Fibroblasts were the predominant cell population (∼62.5%), followed by keratinocytes (∼15%) and immune cells of the myeloid (∼9.4%) and lymphoid (∼7.3%) lineages. Lymphatic (∼3.1%) and vascular (∼1.6%) endothelial cells (EC), that delineate lymphatic and blood vessels, respectively, were also detected, along with vascular smooth muscle cells (SMC) (∼0.5%), which surround larger blood vessels to regulate blood flow. A minor proportion of melanocytes was also present (∼0.4%). These cellular compositions were consistent with prior cutaneous scRNA-seq datasets^46^.

To refine cell annotations, we subclustered the four major cell types based on subtype-enriched genes. Fibroblasts segregated into eight subtypes (Fig. 1G; Fig. S6A, B; Dataset S3), listed in the order of decreasing abundance: *Ccl19^+^* fibroblasts (∼46% of all fibroblasts), which promote tertiary lymphoid structures by recruiting lymphocytes and enhancing local adaptive immune response^47,48^; dermal white adipose tissue (dWAT) fibroblast cells (∼28.2%), which reside beneath the dermis and contribute to wound healing, immune defenses and skin homeostasis^49^; *Pi16^+^* fibroblasts (∼7.3%), which form perivascular sheaths that regulates vascular stability, immune cell extravasation and stromal organization through extracellular matrix (ECM) production and expression of adhesion molecules and chemokines^50–53^; dermal sheath fibroblasts (∼6.4%), which create a contractile niche layer around hair follicles, anchoring them within the skin and participating in dermal repair^54,55^; *Crabp1^+^* fibroblasts (∼4.3%), which modulate retinoic acid signaling, regulating ECM and dermal homeostasis^56,57^; telogen (∼3.1%) and anagen (∼2.5%) dermal papilla (DP) fibroblasts, which constitute the principal hair follicle signaling niche cells and reside at its base, with telogen DP cells maintaining follicle’s quiescence, and anagen DP cells driving active hair growth^58^; and proliferating fibroblasts (∼1.1%), which actively divide to support cutaneous tissue growth, repair and remodeling^59^. Epidermal and ectodermal appendage cells segregated into nine subtypes (Fig. 1H; Fig. S6C, D; Dataset S3): germinative layer cells (∼28.5% of all keratinocytes)^60^, that reside at the base of growing hair follicles where they actively divide, contributing to the steady-state hair growth^61^; basal keratinocytes (∼14.8%), which form the deepest epidermal layer and maintain epidermal attachment^62^; suprabasal keratinocytes (∼12.6%), which form upper layers of the epidermis where they differentiate to form water-tight barrier^61^; sebaceous gland cells (∼11.4%), which produce lipid-rich secretome to lubricate epidermis and hairs, reinforce epidermal barrier and modulate local immune and antibacterial defenses^63^; hair follicle stem cells (HFSCs) (∼8.7%), which reside in the so-called bulge region of the follicle, and critically sustain cyclical hair growth and contribute to epidermal repair after wounding^61^; upper hair follicle keratinocytes (∼8.3%), which form a protective epithelium at the boundary with the epidermis^61^; cortex/cuticle cells (∼8.1%) and medulla cells (∼2.1%), which represent differentiated cell types of the hair shaft; and inner root sheath (IRS) keratinocytes (∼6.6%), which form a protective sleeve around the growing hair shaft^64^. Among myeloid cells (Fig. 1I; Fig. S6E, F; Dataset S3), we identified multiple populations of macrophages: *Spp1*+ macrophages (∼21.6% of myeloid cells), which are associated with sustained inflammation, extracellular matrix and immune response modulation^65^; *Folr2*+ macrophages (∼15.76%), which are perivascular tissue-resident macrophages with the capacity to activate CD8^+^ T cells^66^; monocyte-derived macrophages (moMacro; ∼12.8%), which arise from recruited circulating monocytes and provide a rapidly expandable population that drives inflammatory responses and pathogen clearance^67^ and antigen-presenting macrophages (∼5.95%) with an activated MHC-II program as indicated by *Ciita* high expression^68^. We also observed several DC populations: cDC2 (∼18.0%) and cDC1 (∼5.8%) which activate CD4^+^ and CD8^+^ T cells, respectively^69^; and migratory DCs (∼3.0%) with increased *Ccr7* expression^70,71^. Finally in the myeloid compartment, we identified monocytes (∼10.5%), which have functions in phagocytosis and immune modulation^72^; proliferative myeloids (∼3.5%) with an activated program of cell proliferation (i.e., *Ccna2*, *Spc24*, *Ncapg*, *Ncapd2*, and *Pimreg*); and neutrophils (∼3.2%), which shape early inflammatory response^73^. Among lymphoid cells (Fig. 1J; Fig. S6G, H; Dataset S3), subclustering distinguished: a mixed populations of innate lymphoid cells (ILC), which initiate inflammation through early cytokine secretion^74,75^, and T cells (∼46% of all lymphoid cells); dendritic epidermal T cells (DETC) (∼33.2%), which are skin-resident γδ T cells with roles in immune surveillance and tissue homeostasis^76^; and quiescent-like ILC (∼20.1%), which can differentiate into active ILCs^77,78^. Altogether, these data establish a robust scRNA-seq framework to investigate mosquito bite-initiated viral infection in the skin at single-cell resolution.

### Fibroblasts are the frontline infectious saliva-exposed cells

As mosquito RNA is secreted into the saliva^79,80^, we leveraged it as a molecular proxy to identify skin cell types that directly interact with infectious saliva components. Non-polyadenylated viral reads could not be detected with the scRNAseq technology. Upon data mapping, we aligned scRNA-seq reads against the mosquito genome and identified nearly 300 mosquito-derived reads in bitten skin (after quality controls), whereas none were found in resting control skin. Detected mosquito reads mapped to six genes, two of which—*HISTONE H4* and an uncharacterized locus (*LOC23687614*)—accounted for the large majority (Fig. 2A). Mosquito RNA was detected in 3.9% of all sequenced cells (Fig. 2B), providing an estimate of the limited subset of skin cells directly exposed to mosquito saliva, even with our narrow skin collection method. All annotated skin cell types, except melanocytes, contained mosquito RNA; however, fibroblasts, vascular SMC and keratinocytes exhibited the highest proportion of positive cells (∼ 4%), with fibroblasts being the most exposed in absolute numbers (Fig. 2C). Moreover, only fibroblasts, keratinocytes and myeloid cells contained more than one mosquito read per cell (Fig. S7A-H), supporting their active interaction with mosquito saliva, such as through endocytosis^81^. Intriguingly, among fibroblasts, the distribution of mosquito RNA did not match subtype abundance: proliferating (i.e., the least prevalent subtype) and dWAT fibroblasts exhibited the highest frequencies of mosquito RNA positive cells, followed by *Ccl19^+^* (i.e., the most prevalent subtype) and *Crapb1^+^* fibroblasts (Fig. 2D). This suggests preferential exposure and/or uptake by specific fibroblast subsets.

**Figure 2.**
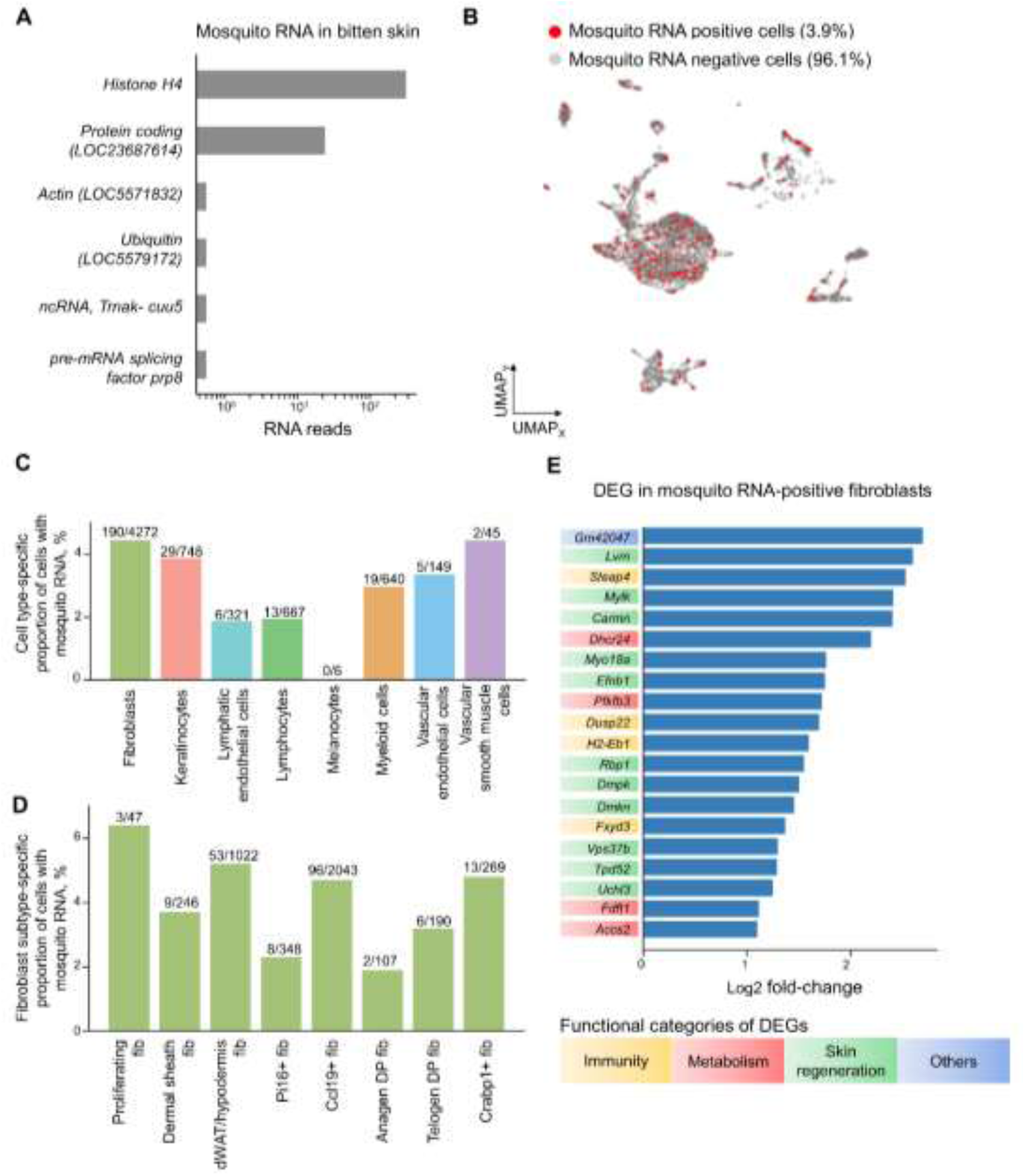
Cellular distribution of mosquito-derived RNAs at the bite site. **(A)** Gene identification of mosquito salivary RNA fragments. **(B)** Distribution of mosquito RNA-positive mouse skin cells. Percent of mosquito RNA-positive cells are indicated in brackets. **(C, D)** Proportion of mosquito RNA-positive cells between cell types (C) and fibroblast subtypes (D). Numbers above the bars indicate absolute number of positive mouse skin cells over total cells. **(E)** Differentially expressed genes (DEG) in mosquito RNA-positive fibroblasts. Functional categories of DEGs are indicated by colored boxes. See also **Figure S7 and Dataset S4.**

Next, we characterized transcriptional response to infectious mosquito saliva by comparing gene expression patterns between mosquito RNA-positive and -negative fibroblasts, identifying 20 significantly upregulated genes based on expression fold-change > 1 (Fig. 2E, Dataset S4). Other non-fibroblast cell types contained too few mosquito RNA-positive cells to support statistically robust analyses. In fibroblasts, four differentially-expressed genes (DEGs) are associated with immune-related functions: *Fxyd3* enhances Il-17A signalling, promoting cutaneous inflammation^82^; *H2-Eb1* encodes an MHC-II protein that drives adaptive immune activation^83^; *Dusp22* modulates T cell activation through the JNK pathway^84^; and *Steap4* responds to inflammation-induced oxidative stress^85^. Four DEGs are related to metabolic pathways, inducing steroid biosynthesis (*Dhcr24*^86^ and *Fdft1*^87^) and energy-producing glycolysis (*Acss2*^88^ and *Pfkfb3*^89^). Three DEGs are associated with skin repair and tissue remodeling: *Dmkn* promotes cutaneous barrier formation^90^; *Efnb1* is an adhesion molecule with an established role in promoting angiogenesis^91^; and *Dmpk* drives myoblast differentiation^92^. Three DEGs promote cell motility: *Mylk* and *Myo18A* alter cytoskeleton^93,94^, and *Uchl3* influences epithelial-mesenchymal transition^95^. Five DEGs are involved in general cellular homeostasis: *Lvrn* is implicated in ECM remodeling^96^; *Tpd52* and *Rbp1* in promoting cell proliferation^97,98^; *Carmn*, a long non-coding RNA, in regulating cell differentiation^99^; and *Vps37B* in endosomal sorting through the ESCRT pathway^100^. Collectively, these scRNA-seq analyses provide the first single-cell-level identification of cutaneous cell types exposed to mosquito saliva and highlight fibroblasts, a structural cell type, as primary responders during the earliest stages of vector-host interactions.

### Myeloid and lymphoid immune cells are remodeled at the bite site

To characterize cutaneous cellular dynamics in response to infectious bites, we compared cellular composition of bitten *vs.* resting skin in our scRNA-seq dataset via permutation testing. At the cell type level, we observed a reduction in melanocyte and keratinocyte populations, whereas lymphocytes, vascular SMCs, vascular and lymphatic ECs expanded following infectious bites (Fig. 3A-C). Among fibroblast subtypes, telogen DP cells were markedly enriched upon infectious bite (Fig. 3D-F). Among keratinocytes and ectodermal appendage cells, populations of germinative layer, medulla, cortex and IRS strongly contracted, while sebaceous and suprabasal keratinocyte populations expanded (Fig. 3G-I). Among myeloid cells, abundances of Folr2 macrophages and neutrophils were reduced, whereas monocyte-derived and antigen-presenting macrophage populations were strongly increased (Fig. 3J-L). Additionally, monocyte and proliferative myeloid compartments were marginally enhanced from 8.2 to 12.5% and from 2.8 to 4%, respectively. Interestingly, population sizes of migratory DCs remained unchanged. Among lymphocytes, quiescent-like ILCs and DETC were strongly recruited to the bite site, increasing by about 4-fold (Fig. 3M- O). Collectively, while the observed hair follicle cell population differences might reflect natural hair cycle heterogeneity between animals^101^, differences in the other cell types reveal rapid and extensive multicellular remodeling of the skin in response to an infectious bite, establishing the early recruitment of multiple types of innate and adaptive immune cells.

**Figure 3.**
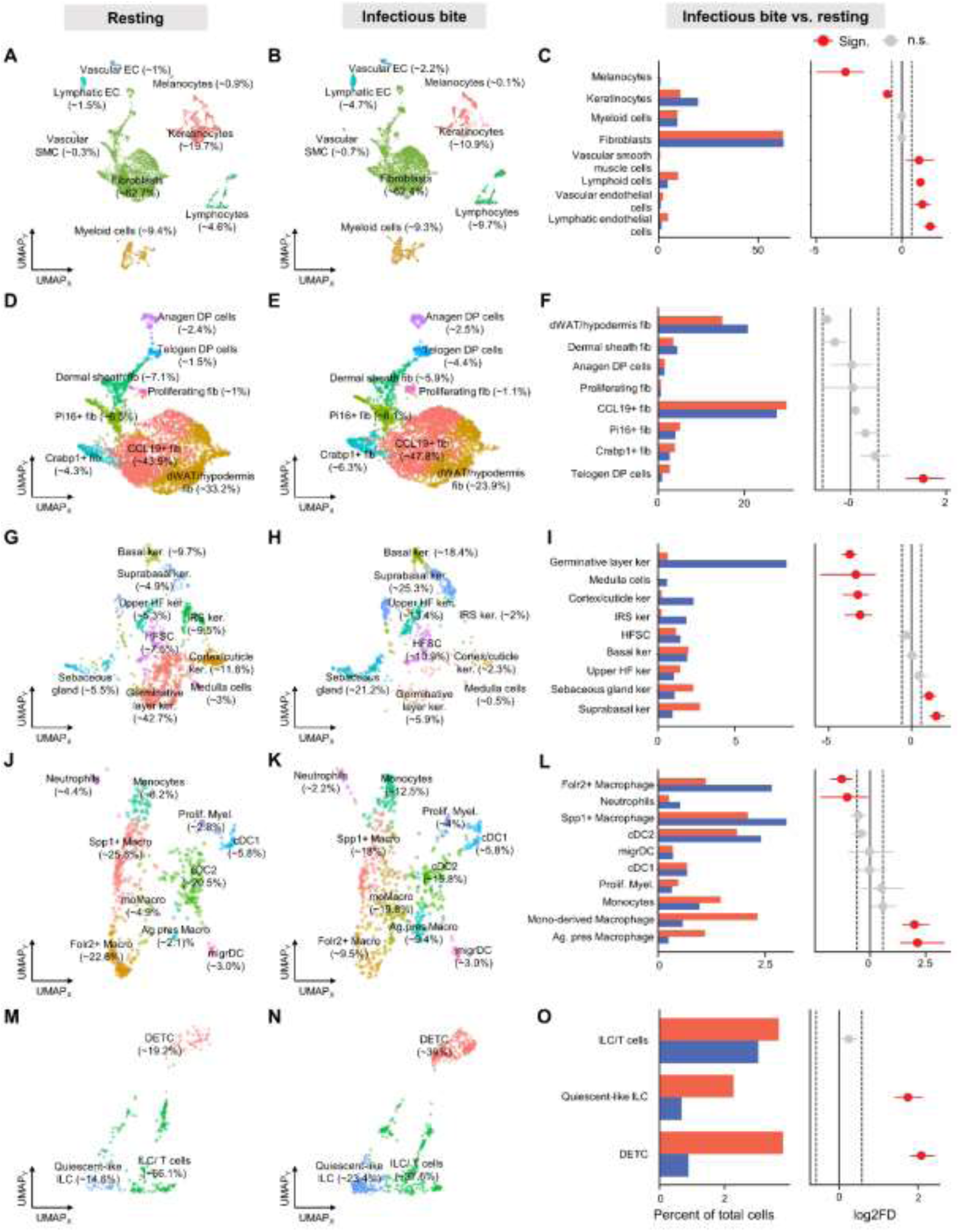
Cellular landscape of mouse skin after infectious bite. **(A-C)** Distribution of cell types in unbitten skin (resting) (A) and in skin at 6 hours post-infectious bite (infectious bite) (B), and percent comparisons and cell type fold change between infectious *vs.* rest conditions (C). **(D-O)** Distribution of cell subtypes among fibroblasts (D-F), keratinocytes and ectodermal appendage cells (G-I), myeloid cells (J-L), and lymphocytes (M-O) in resting skin and skin at 6 hours post-infectious bite. (A, B) Proportions within all cells are shown in brackets. (D, E, G, H, J, K, M, N) Within-cell type proportions are shown in brackets. (C, F, I, L, O) Changes in cell proportions are expressed as log2 fold difference (Log2FD) means ± standard deviation in infectious bite as compared to resting conditions. Dashed black lines indicate a fold change of 1.5. Statistical significance indicated in red for p < 0.05 based on permutation test. ECs, endothelial cells; SMCs, smooth muscle cells; DP, dermal papilla; dWAT, dermal white adipose tissue; HF, hair follicle; IRS, inner root sheath; DCs, dendritic cells; HFSC, hair follicle stem cells; DETC, dendritic epidermal T cells; ILC, innate lymphoid T cells.

### Fibroblasts and myeloid cells orchestrate immune signaling response to infectious bite

To investigate how an infectious bite reshapes microenvironmental signaling in the skin, we inferred cell-cell communication (CCC) activity between cell subtypes from the scRNA-seq dataset using CellChat^102,103^. CellChat database annotates ligands, receptors, and signaling agonists/antagonists across CCC pathways. First, we assessed global shifts in CCC of the skin by aggregating all ligand-receptor interactions significantly present in the data (Fig. 4A). This overview revealed that infectious bite enhanced communication between fibroblast subtypes, while reducing signaling from fibroblasts to keratinocytes and between keratinocyte subtypes. Among fibroblasts, *Crabp1^+^* cells exhibited the strongest activation, both as signal senders and receivers. Among immune cells, communication across myeloid subtypes was moderately increased, especially between neutrophils, monocytes and macrophages. Next, we quantified the activity of individual CCC pathways. Of 62 pathways that were detected as active in either sample, 24 were upregulated and 11 downregulated by infectious bite (Fig. 4B). Notably, upregulated pathways were predominantly immune-related (18 out 24), whereas downregulated pathways were mostly associated with skin repair (7 out of 11).

**Figure 4.**
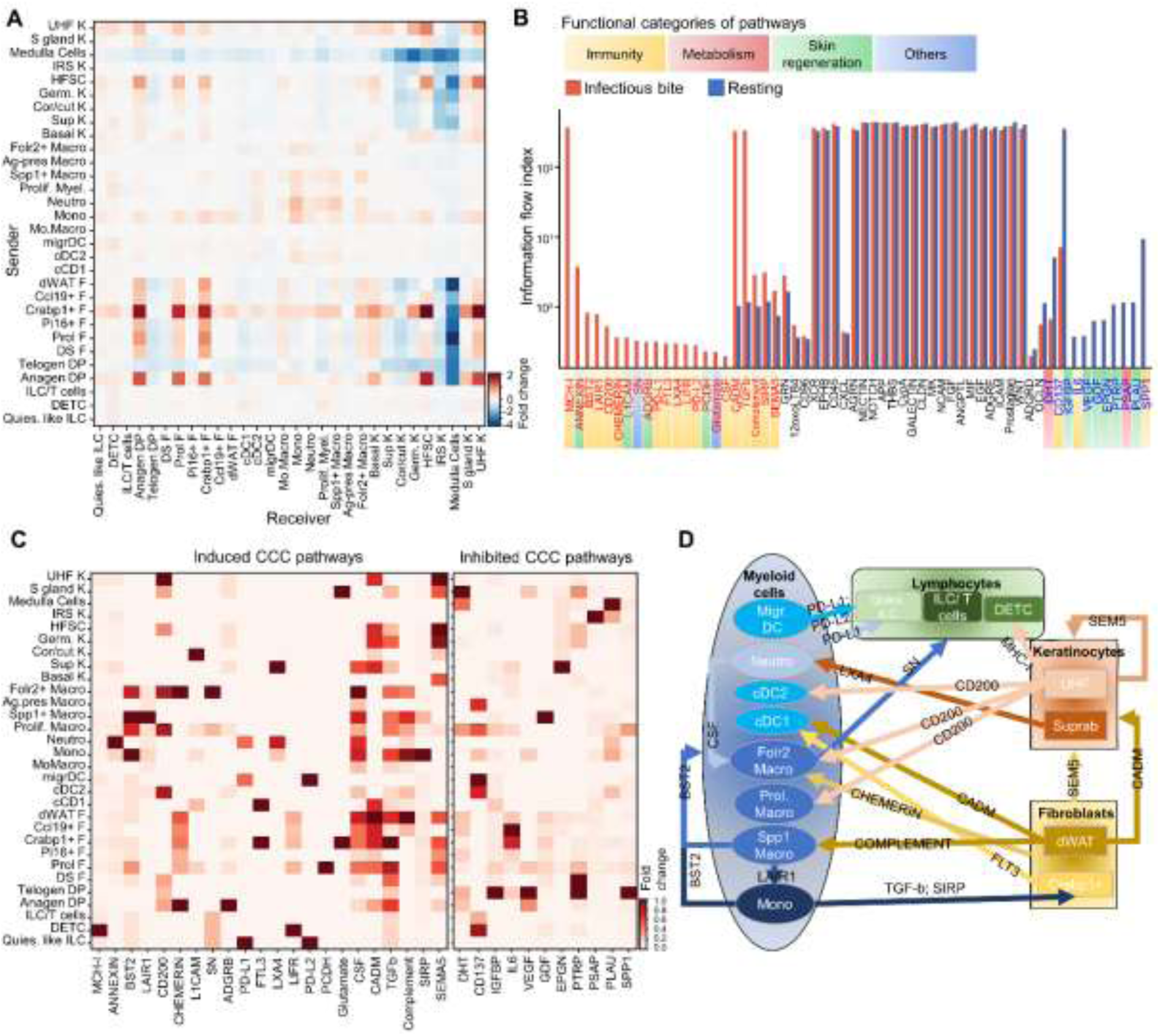
Cell-cell communication changes in response to infectious bite. **(A)** Influence of infectious bite on the overall cell-cell communication (CCC) strength between skin cell subtypes. NK, natural killer; F, fibroblasts; DP, dermal papilla; DS, dermal sheath; migr, migratory; Prol., proliferative; Macro, macrophage; Neutro, neutrophil; mono, monocytes; K, keratinocytes; Sup, suprabasal; Cor/cut, cortex/cuticle; Germ, germinative; HFSC, hair follicle stem cells; IRS, inner root sheath; S, sebaceous; UHF, upper hair follicle. **(B)** Pathway information flow among active CCC pathways in resting and infectious bite conditions. Pathway names in red indicate significant induction and in blue significant inhibition by infectious bite (p < 0.001; as determined by Wilcoxon test^102^). Functional categories of regulated pathways are indicated by colored boxes. **(C)** Influence of infectious bite on signaling strength for the significantly induced and inhibited pathways per cell subtype. **(D)** Overview of the induced CCC immune pathways between cell subtypes. See also **Figure S8.**

We mapped the activity of regulated pathways to specific cell subtypes (Fig. 4C) and detailed their contributions as signal senders, receivers, mediators and/or influencers (Fig. S8). Focusing on immunity, we integrated the upregulated interactions to delineate global immune CCC patterns (Fig. 4E). Fibroblasts emerged as active communicators, engaging six CCC pathways toward keratinocytes and myeloid cells. Toward keratinocytes, most fibroblast subtypes induced pro-inflammatory SEMA5 signaling^104^, while dWAT subtypes triggered CADM-mediated immune cell adhesion^105^. Chemotactic CHEMERIN signaling^106^ was broadly promoted in *Folr2*^+^ macrophages by multiple fibroblast subtypes. In contrast, dWAT and *Crabp1^+^* fibroblasts activated pro-inflammatory CADM^107^ and hematopoietic-activating FLT3^108^ signaling, respectively, in cDC1. Additionally, dWAT fibroblasts induced COMPLEMENT signaling in *Spp1*^+^ macrophages. Keratinocytes coordinated three CCC pathways, directing signals toward keratinocytes, myeloid cells and NK cells. Reciprocal SEMA5-mediated inflammatory signaling occurred between keratinocyte subtypes, while upper hair follicle keratinocytes stimulated MHC-I-dependent antigen-presentation^109^ in DETC. In contrast, suprabasal keratinocytes promoted anti-inflammatory LXA4 signaling in neutrophils^110^ and upper hair follicle induced anti-inflammatory CD200^111^ in cDC2, *Folr2*^+^ and proliferative macrophages. The infectious bite provoked the most extensive reorganization of intercellular communication within the myeloid compartment, activating eight distinct pathways targeting fibroblasts, myeloid subtypes and lymphocytes. Monocytes induced both immuno-modulatory TGFβ signaling^112^ and immuno-inhibitory SIRP signaling^113^ in *Crabp1^+^* fibroblasts, while cooperatively stimulating anti-viral BST2 signaling^114^ with *Spp1*^+^ macrophages across myeloid subtypes. In return, *Spp1*^+^ macrophages promoted anti-inflammatory LAIR1 signaling^115^ in monocytes, highlighting layered regulatory mechanisms. Neutrophils stimulated CSF-dependent phagocytosis^116^ in *Folr2*^+^ macrophages, which induced phagocytosis-promoting SN signaling^117^ across lymphocyte populations. Conversely, neutrophils and migratory DCs co-activated immuno-modulatory PD-L1 and PD-L2 pathways^118^ in quiescent-like ILCs. Taken together, these CellChat analyses illustrate how infectious bite rapidly reprograms the cutaneous signaling networks, dampening tissue repair pathways while amplifying pro- and anti-immune-directed communication across stromal, epithelial and immune compartments.

### Cell subtype-specific responses to infectious bite integrate immune, skin repair and metabolic transcriptional programs

To assess cell subtype-specific responses to infectious bite, we identified DEGs between infectious and resting conditions within each cell subtype using a conservative fold-change threshold of ≥ 4, revealing 2,953 DEGs (Fig. 5A; Dataset S5). Keratinocytes and ectodermal appendages showed the most extensive transcriptional reprogramming (986 DEGs: 699 up, 287 down), with basal, cortex/cuticle, germinative layer, and IRS cells exhibiting the largest responses (>162 DEGs each), whereas HFSCs, medulla, sebaceous gland, and upper hair follicle cells had more modest responses (<66 DEGs each). Skin repair-associated genes were predominantly regulated, followed by immune-related genes, which accounted for more than 23.3% of DEGs in cortex/cuticle, germinative, HFSC, and IRS cells, with metabolic genes also being significantly modulated (Fig. S9A). Fibroblast subtypes featured fewer overall DEGs (717 DEGs: 289 up, 428 down) (Fig. 5A; Dataset S5), albeit *Ccl19^+^, Crabp1^+^,* dermal sheath, dWAT, and *Pi16^+^* fibroblasts each had between 94 and 124 DEGs. Skin repair-associated genes were most regulated, followed by immune- and metabolic-related genes (Fig. S9A). Among myeloid cells featuring 745 DEGs (193 up, 552 down) (Fig. 5A; Dataset S5), moMacrophages were the most responsive (274 DEGs), followed by antigen-presenting macrophages (148 DEGs), while other subtypes exhibited a moderate response (<61). Interestingly, while skin repair-associated genes were also highly regulated across myeloid cell subtypes, immune-related genes were the most regulated functional category in migratory DCs and present in other subtypes (Fig. S9A). Metabolic genes were modulated across all myeloid cell subtypes. Among lymphocytes, DEGs were predominantly downregulated (353 DEGs: 78 up, 275 down) and fairly homogeneous across DETCs, quiescent-like ILCs and ILC/T cells (125, 122, and 106 DEGs, respectively) (Fig. 5A; Dataset S5). Here too, skin repair-associated genes were predominant, followed by immune- and metabolic-related genes (Fig. S9A). Finally, lymphatic and vascular ECs and SMCs showed moderate responses (76, 54, and 22 DEGs, respectively) (Fig. 5A; Dataset S5), mainly involving skin repair, with minor immune- and metabolism-related regulation.

**Figure 5.**
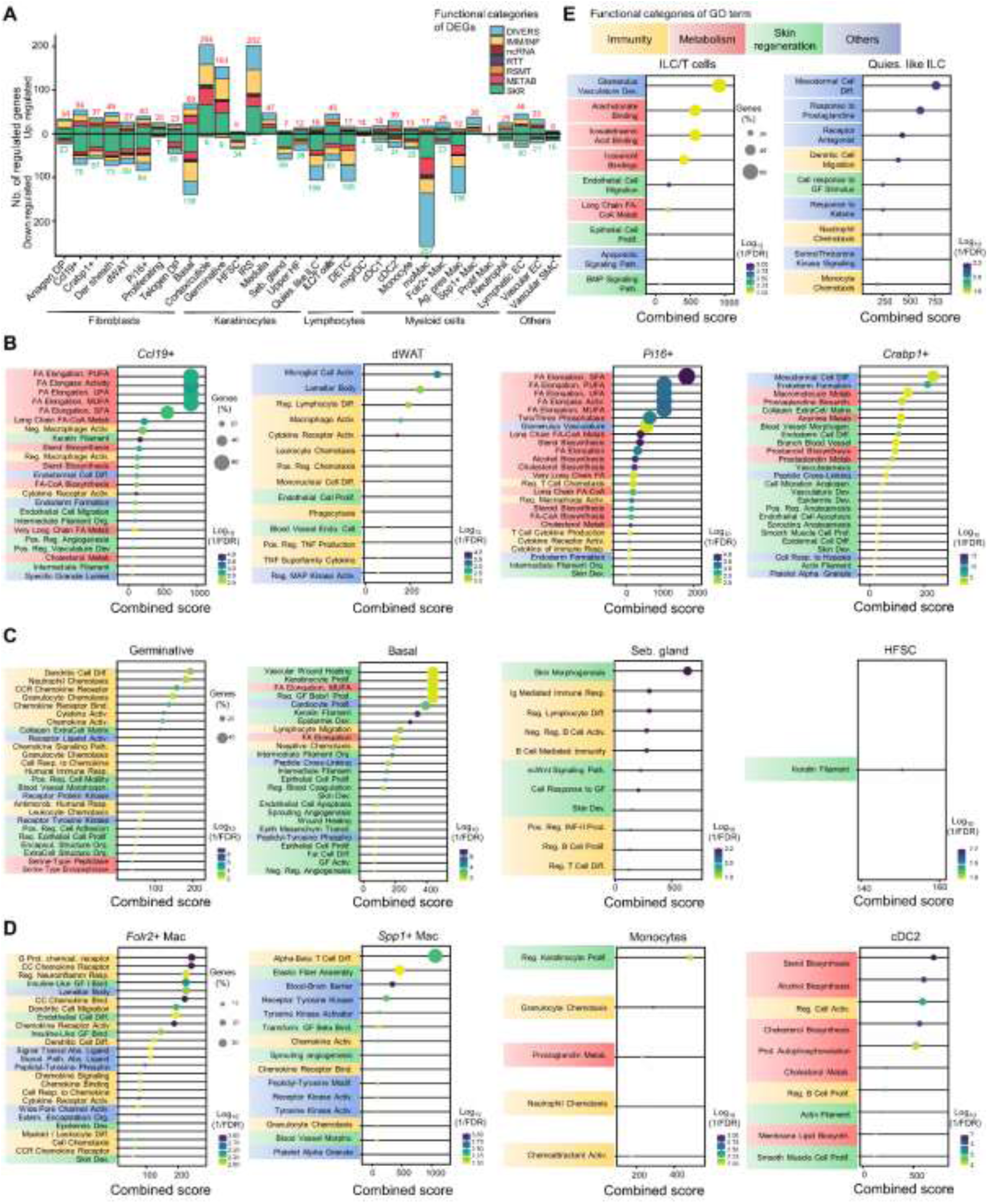
Cell-specific transcriptomic response to infectious bite. **(A)** Functional distribution of DEGs (adj. p-value < 0.001; |fold change| ≥ 4) within cell subtypes. Bars indicate the numbers of regulated genes. Functional categories: SKR, Skin Regeneration; METAB, Metabolism; RSMT, Redox-Stress-Mitochondria; RTT, replication, transcription and translation; ncRNA, non-coding RNA; IMM/INF, immunity and inflammation; DIVERS, other and unknown functions. N total DEGs, 2,481. Number of up- and down-regulated DEGs are indicated in red and green, respectively. (**B-E**) Gene Ontology (GO) biological process enrichment analysis for the top 4 most abundant subtypes of fibroblasts (B), keratinocytes and ectodermal appendage cells (C), myeloid cells (D) and lymphocytes (E). Subtypes are presented in an increasing order of cell abundance. Significant GO terms encompass > 10% all DEGs within subtypes and have p-value < 0.01. Functional categories are indicated by colored boxes. dWAT, dermal white adipose tissue cells; IRS, inner root sheath cells; DC, dendritic cells; DETC, dendritic epidermal T cells; ILC, innate lymphoid cells. See also **Figure S9 and Dataset S5.**

To refine characterization of transcriptional remodeling across cell subtypes, we performed GO analyses on within-subtype upregulated DEGs using stringent thresholds (>10% DEGs; *p* < 0.01), presenting results in a decreasing order of cell abundance. Among fibroblasts (Fig. 5B), *Ccl19*⁺ cells showed robust activation of fatty-acid biosynthesis, accompanied by modulation of macrophage activation and induction of the pro-angiogenic program. dWAT fibroblasts predominantly upregulated immune-related processes, including several lymphoid-targeting chemoattractants and phagocytosis-related genes. *Pi16⁺* and dermal sheath fibroblasts exhibited strong fatty-acid biosynthetic signatures and moderate chemotactic activity—directed toward T cells and macrophages in *Pi16⁺* cells, or toward leukocytes in dermal sheath cells—and both contributed modestly to skin repair programs. In contrast, *Crabp1⁺* fibroblasts strongly induced skin repair pathways, particularly those linked to angiogenesis and dermal development, while moderately upregulating prostaglandin biosynthesis genes. Anagen DP fibroblasts preferentially activated T cell-related pathways, whereas telogen DP fibroblasts elicited a more modest neutrophil-attracting signature (Fig. S9B). Proliferating fibroblasts uniquely upregulated endothelial development pathway genes.

Among keratinocytes and ectodermal appendage cells (Fig. 5C), germinative cells markedly elicited granulocyte and neutrophil chemoattraction and antimicrobial humoral responses, while also contributing to ECM-driven and cell migratory processes associated with skin repair, but with minimal engagement of metabolic pathways. Basal keratinocytes predominantly activated skin-repair programs, including wound healing and skin development pathways, while minimally modulating immune or metabolic functions. Sebaceous gland cells displayed an immune-centric response focused on B-cell activation, together with the induction of tissue growth associated pathways. Both cortex/cuticle cells and IRS primarily upregulated skin repair programs, with moderate contributions to immune chemoattraction (Fig. S9C). HFSCs and medulla cells each displayed only a single enriched biological process.

Among myeloid cells (Fig. 5D), *Folr2*^+^ macrophages exhibited strong activation of chemoattraction-related GO terms, accompanied by a moderate induction of myeloid differentiation and skin development. *Spp1*^+^ macrophages contributed to T cell differentiation, chemotaxis and angiogenesis. Monocytes showed enrichment of GO terms related to myeloid chemoattraction, keratinocyte proliferation and prostaglandin metabolism. cDC2 had major involvement in sterol-related metabolism, with a modest implication in B cell activation. Neutrophils uniquely displayed T cell receptor complex enrichment (Fig. S9D). Among lymphocytes (Fig. 5E), whereas DETC cells lacked significant GO enrichment, ILC/T cells did not upregulate classical immune-response pathways but, instead, prominently activated eicosanoid biosynthesis genes. Quiescent-like ILCs, in contrast, activated chemotactic programs targeting DCs, neutrophils, and monocytes, and contributed moderately to skin development processes.

Finally, lymphatic ECs engaged a mix of immune, metabolic, and tissue repair responses; vascular ECs displayed modest immune-related signature; and vascular SMCs showed limited immune- and tissue repair-related activation (Fig. S9E-G). Together, these GO analyzes indicate that infectious mosquito bites trigger a multifaceted, cell subtype-specific transcriptional response integrating skin repair, immune activation, and metabolic adaptation across diverse tissue compartments, revealing specialized functions for distinct cell populations.

### Fibroblast response to infectious bite reduces viral transmission

To determine how the above described cutaneous transcriptional responses contribute to viral transmission, we selected three antiviral DEGs in fibroblasts; *Saa2* primarily upregulated in Pi16+ fibroblasts and three other subtypes induces anti-viral IFN response^119^, *Defb8* upregulated in DP*, Ccl19^+^* and dWAT fibroblasts directly acts on viruses and *Lrrc15* selectively induced in Crabp1+ fibroblasts has demonstrated function against viruses^120,121^ (Fig. 6A; Fig. S10A-C). We first assessed their regulation following *in vitro* exposure to virus, saliva (mimicking the bite), or both in the lineage-matched human cell models: human foreskin fibroblasts (HFF1) (Fig. 6B). Control experiments confirmed that the dye used for saliva collection^44^ and saliva supplementation alone did not alter cell viability or infection rates (Fig. S10B, C). At 6 hours post-infection, only *LRRC15* was modulated by virus in a manner concordant with the *in vivo* regulation in mouse skin (Fig. 6C), whereas the remaining genes displayed inverse patterns (Fig. S10D). Next, we depleted *LRRC15* in HFF1 to interrogate its function on WNV infection (Fig. 6D). Notably, siRNA-mediated reduction of LRRC15 mRNA and protein levels (Fig. S10E, F) triggered a strong viral enhancement (Fig. 6E), identifying LRRC15 as a potent restriction factor.

**Figure 6.**
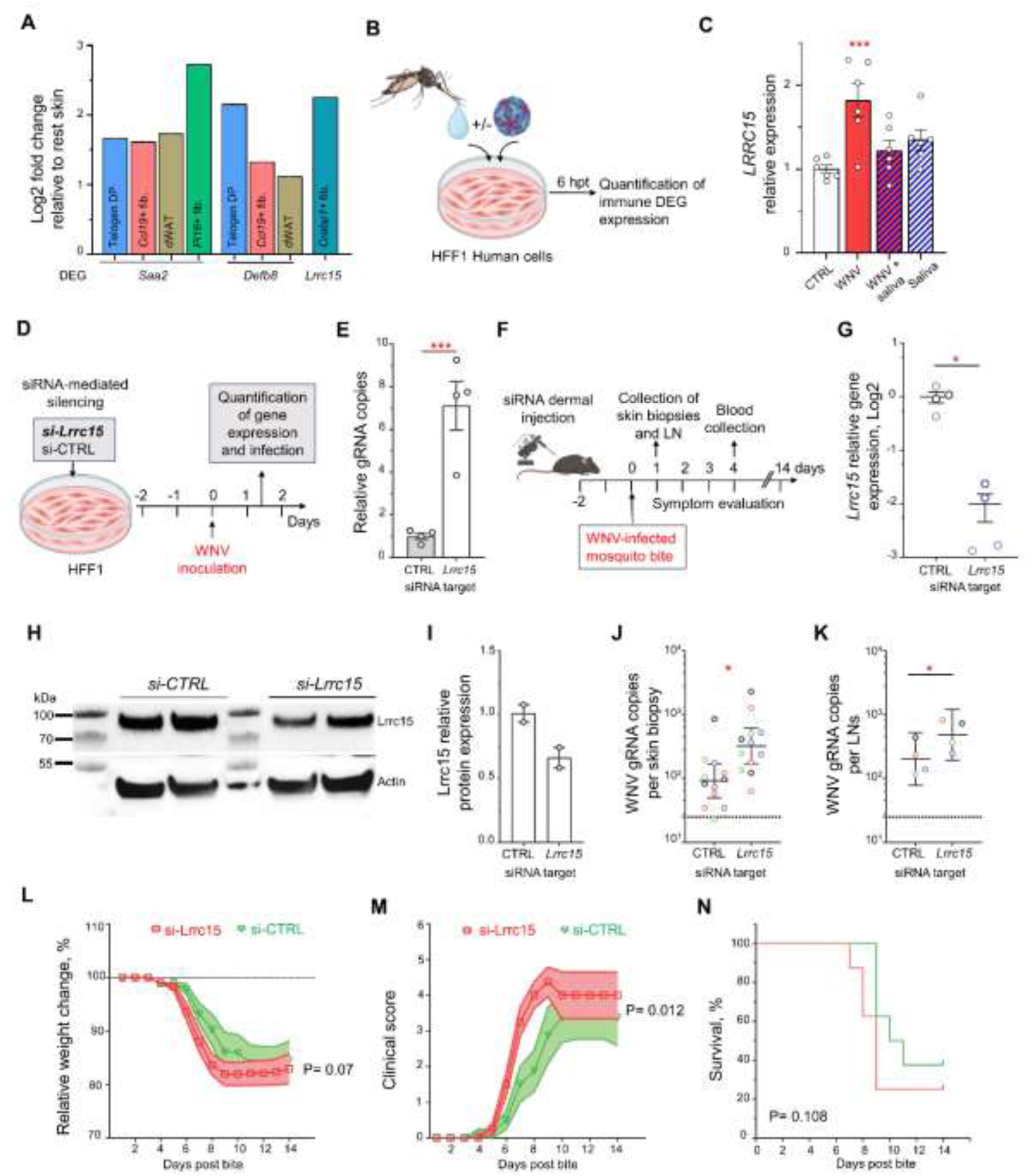
Fibroblast response to infectious bite reduces transmission. Expression of selected immune DEGs in fibroblasts. **(A)** Infection of dermal fibroblast (HFF1) cells with or without mosquito saliva and quantification of selected DEGs. hpt, hours post treatment. **(B)** Regulation of *LRRC15* expression by either saliva, virus (WNV) or both in HFF1. CTRL, mock infection. DEGs that were not regulated as in skin scRNA-seq dataset are presented in Figure S10. **(C)** Functional evaluation on WNV infection of LRRC15 in HFF1. CTRL, siRNA control. **(D)** Infection levels in *LRRC15*-depleted HFF1 cells at 36 hours (h) post-infection. **(E)** Evaluation of Lrrc15 function in transmission. **(G-I)** Expression of Lrrc15 gene (G) and protein (H,I) in mouse skin at 48 h post-siRNA injection. N mice, 2. Each lane represents the sample from one mouse. **(J, K)** Infection level per skin biopsy (J) and pair of inguinal LNs (K) at 24 h post-infectious bite. N mice, 4. Skin biopsies and LN from the same mouse are color-coded. **(L-N)** Relative weight changes (L), clinical scores (M) and survival (N) over 14 days post-infectious bite in mice intradermally injected with siRNA against *Lrrc15* or control siRNA. N mice, 8. (C, E, G, I-K) Bars and lines show means ± s.e.m. Dots indicate repeats. *, p < 0.05; **, p < 0.01; ***, p < 0.001 by Dunett’s test with CTRL or T-test. (L, M) Lines indicate mean and shaded area ± s.e.m. p, as determined by mixed-effect ANOVA for time x treatment interaction (L) and treatment effect (M). (N) p, as determined by Kaplan-Meier Survival test. See also **Figures S10-S11 and Datasets S1, S5.**

Next, we developed an *in vivo* cutaneous siRNA-mediated silencing approach in mice to investigate the effect of *Lrrc15* on transmission (Fig. 6F). Forty-eight hours after siRNA injection, when cutaneous levels of *Lrrc15* transcript (Fig. 6G) and protein (Fig. 6H, I) were reduced, skin site was exposed to infectious bites. Local Lrrc15 depletion enhanced viral load in both skin (Fig. 6J) and draining LNs (Fig. 6K) at 24 hpb. Furthermore, while RNAemia at 4 days post-bite confirmed transmission (Fig. S10G), *Lrrc15* silencing exacerbated weight loss (Fig. 6L), disease severity (Fig. 6M; Dataset S1) and animal mortality (Fig. 6N). Collectively, these findings demonstrate that fibroblast-expressed *Lrrc15* functions as a critical antiviral factor restraining early mosquito-borne infection and underscore the essential contribution of dermal fibroblast responses to arbovirus transmission.

## Discussion

The initial cutaneous response to an infectious mosquito bite is a critical determinant of arbovirus transmission success^41^. To dissect the cellular and molecular architecture of this decisive early event, we developed a scRNA-seq protocol targeting a minimal skin area at the site of WNV-infected mosquito bites in immunocompetent mice, precisely during the transmission bottleneck phase. Through integrated bioinformatic analyses, we defined the skin cell composition, identified mosquito saliva-exposed cells, inferred cell-cell communication networks, and characterized cell type-specific transcriptional programs, providing a high-resolution map of the earliest stages of arbovirus transmission. These analyses reveal distinct, subtype-specific cellular roles and underscore the essential contribution of non-immune, structural skin cells—particularly fibroblasts—to infection establishment. Using skin-localized gene depletion, we demonstrate that fibroblast-intrinsic responses are key determinants of transmission efficiency and downstream pathogenesis. By combining high-resolution transcriptional profiling with functional perturbation approaches, this study provides an integrated, multiscale view of the cross-cellular response to an infectious mosquito bite, illuminating the molecular and cellular dynamics that govern arbovirus transmission.

The orchestration of myeloid cell response in the skin is a critical driver of arbovirus transmission^41,122^. This response is first initiated by a rapid influx of neutrophils, detectable as early as 3 hours after an uninfectious bite in mice and 4 hours in humans, followed by a decline in neutrophil abundance by 5^th^ hour post-bite in mice^17,19^. Consistent with this kinetic profile, we observed a reduced neutrophil population at 6 hours post-infectious bite, indicative of ongoing resolution of neutrophil-driven inflammation^123,124^, further supported by the anti-inflammatory LXA4 signalling^110^ toward neutrophils. Rapid attenuation of non-infectious bite-induced inflammation has similarly been described in human skin through the differentiation of anti-inflammatory M2-like macrophages^19,125^. In our study, *Folr2*^+^ macrophage abundance was reduced following infectious bite, potentially resulting from inflammation resolution^126^. Despite these resolution-associated signatures, inflammation remained active at 6 hours post-infectious bite, as reflected by sustained TNF expression in neutrophils, multiple chemokine expressions in cDC2, *Folr2*^+^ macrophages and *Spp1*^+^ macrophages (Dataset S5), enrichment of GO terms associated with chemoattraction, and pro-inflammatory signaling. We also observed a second wave of myeloid cell recruitment^19,41^, comprising monocyte-derived, antigen-presenting macrophages, monocytes and proliferative myeloids, although at non- significant levels for the two latter. While all myeloid populations displayed a mixed transcriptional response to infectious bite characterized by concurrent up- and down-regulation of cytokines, coexisting pro- and anti-inflammatory signaling pathways, and GO terms linked to leukocyte chemoattraction, migratory DCs had a clear signature of induced anti-viral interferon response (IFIT1, IRF7, ISG15) (Dataset S5). Our results detail with molecular resolution the dynamic and balanced nature of myeloid responses at early stages of bite-initiated transmission, delineating a tightly regulated cutaneous innate immune landscape, in which inflammatory activation and resolution programs are simultaneously engaged.

Within the lymphoid compartment, we observed a robust influx of DETCs, skin-resident sentinels that mount early defense responses through rapid release of inflammatory mediators upon activation ^76^. Consistent with a role in viral transmission, diverse viral infections similarly recruit DETCs, which subsequently orchestrate myeloid-cell infiltration^127^ and modulate early infection dynamics^128^. Infectious mosquito bite also expanded a population of quiescent-like ILCs, suggesting the formation of a poised reservoir of tissue-resident precursors capable of rapidly differentiating into effector ILCs upon exposure to inflammatory or viral cues^77,78^. Alternatively, their accumulation may reflect active immunomodulation by saliva components, as indicated by the co-induction of anti-inflammatory PD-L1 and PD-L2 pathways^118^. Despite an overall bias toward transcriptional restraint, both DETCs and quiescent-like ILCs remained responsive to infectious bite, showing bidirectional regulation of inflammatory chemokines, interleukins, and immunoregulatory pathways. In contrast, the mixed ILC/T-cell compartment exhibited stronger transcriptional activation and, along with other lymphocyte subsets, was preferentially targeted by SN, which promotes phagocytic interactions^117^. Collectively, extensive recruitment and activation of lymphoid cells at the bite site indicate their potential and previously underappreciated role in shaping early events that influence viral transmission.

Our untargeted approach details the contribution of structural cells, which have the capacity to mount an innate immune response^59,129^, in actively shaping the broader cutaneous immune response to an infectious bite. All fibroblast and keratinocyte subtypes showed enrichment for inflammatory genes and chemoattraction pathways (Dataset S5), with dWAT fibroblasts and germinative keratinocytes exhibiting a particularly strong immune-associated transcriptional profile. Structural cells also displayed direct anti-viral features including defensins^130^ in *Ccl19^+^* and DP fibroblasts, and in basal, germinative, IRS, cortex/cuticle keratinocytes; cytotoxic mediators (granzymes and perforin) in dWAT and *Pi16*⁺ fibroblasts, and in germinative keratinocytes; pattern-recognition receptors (C-type lectins) in germinative keratinocytes; and multiple interferon-inducible genes (Dataset S5). CCC analyses suggest that fibroblasts and keratinocytes engage with myeloid cells, activating pathways linked to pathogen control (CADM^107^, MHC-I^109^, complement^131^), immune cell proliferation (FLT3^132^), recruitment (chemerin^106^, SEMA5^133^), and regulation of inflammation (LXA4^134^, CD200^135^). Together, these data indicate that structural skin cells adopt immune-active states following infectious bite, contributing to early innate defense and immune orchestration.

A major finding of our study is the *in vivo* identification of an anti-transmission response mediated by fibroblasts, revealing fibroblast-expressed LRRC15 as a potent restricting factor for WNV. Being predominantly expressed in the skin^120,136^, *LRRC15* encodes a membrane protein related to Toll-like receptors^120^, a family of pattern-recognition receptors that initiate innate immune signaling^137,138^. Consistent with this function, LRRC15 induces a robust antiviral program in fibroblasts and limits infection by SARS-CoV-2 and adenovirus^120,121^, supporting a broad anti-viral function in line with our findings. Using localized depletion of Lrrc15 in mouse skin, we demonstrate that Lrrc15 restricts viral transmission and attenuates disease severity. Underscoring its potential relevance to viral transmission in humans, meta-analysis of single-cell skin data part of Human Cell Atlas (HCA) project, which consists of 463,503 cells integrated from 24 published studies, revealed prominent *LRRC15* expression primarily restricted to ‘secretory’ and ‘mesenchymal’ fibroblast types enriched in upper dermis and around hair follicles, respectively and present across multiple body locations (Fig. S11A-I). A central unresolved question in vector-borne viral transmission concerns the identity of the initial target cells following exposure to infectious saliva. Our results demonstrate that productive infection of fibroblasts is required for transmission and, together with their pronounced exposure to bite-delivered infectious saliva, support the conclusion that fibroblasts constitute the initial cellular targets. Although virus is detectable in draining LNs as early as 10 minutes post-bite, consistent with reported rapid lymphatic viral dissemination^25,139^, restricting local infection in skin fibroblasts markedly reduces LN seeding and disease outcome, establishing fibroblasts as critical regulators of early viral spread and transmission.

### Limitations of the study

Despite numerous insights that our study offers, it has several limitations. First, our analysis provides a snapshot of the skin response at 6 hpb, capturing early dynamics but not the full temporal trajectory of infection, immune activation, and tissue remodeling. Second, although mice provide a tractable and highly susceptible model for WNV, inter-species differences in skin architecture and immune composition may limit direct translation to humans, albeit prominent fibroblast-specific expression of *LRRC15* in skin HCA dataset suggests translatability. Similarly, our study was conducted with one mosquito vector species and require further experimentation to expand our findings to other vector species. Finally, as our work centers on WNV, it remains uncertain whether the identified fibroblast responses, including the restricting function of LRRC15, represent a generalizable mechanism across arboviruses.

## Material and methods

### Cells

African green monkey kidney Vero (CCL-81), HFF-1 (SCRC-1041), U937 (CRL-1593.2), and Jurkat (TIB-152) cells were obtained from ATCC. Vero and HFF-1 cells were cultured in Dulbecco’s Modified Eagle Medium (DMEM) (Gibco). U937 and Jurkat were maintained in Roswell Park Memorial Institute (RPMI) (Gibco) medium. All media were supplemented with 10% heat-inactivated fetal bovine serum (FBS) (Eurobio), except for HFF-1 which were grown with 15% FBS, and 1% penicillin/streptomycin mix (Invitrogen). Cells were incubated at 37°C with 5% CO2. All cell lines were tested annually for mycoplasma contamination using specific primers and consistently tested negative.

### Viruses

West Nile virus (WNV) strain IS98-ST1 (or Stork 98) was isolated from a stork in Israel in 1998 ^140^ and obtained from Dr. Philippe Desprès, Centre de Ressources Biologiques, Institut Pasteur, Paris. Virus was propagated in C6/36 cells with 2% FBS, titrated using plaque assay with Vero cells as described previously ^141^, and stored at - 70°C.

### Mice

Six- to eight-week-old male C57BL/6J mice were obtained from Charles River (France) and housed in ventilated NexGen Mouse 500 cages (Allentown; Serial number: 1304A0078) within the biosafety level 3 (BSL-3) animal facility at MIVEGEC-IRD, Montpellier, France. They were maintained under a 17 h:7 h light/dark cycle at 53–57% humidity and a temperature of 20–24°C, with *ad libitum* access to sterilized water and an irradiation-sterilized mouse diet (A03, SAFE, France). To allow acclimatization, mice were used for experiments one week after arrival. All efforts were made to minimize pain and stress. Animal protocols were approved by the APAFiS national ethical committee (permission numbers: DAP31273, DAP43466).

### Mosquitoes

The *Aedes aegypti* BORA colony, originally collected from Bora-Bora Island in 1980 ^142^, was maintained at the VectoPole insectary, MIVEGEC. Eggs were hatched in deionized water, and larvae were reared at 26°C under a 12 h:12 h light-dark cycle, receiving ground fish food (TetraMin, Tetra) until pupation. Adult mosquitoes were housed in Bioquip cages at 28°C with 70% relative humidity, a 14h:10h light-dark cycle, and continuous access to a 10% sugar water solution.

### Mosquito inoculation

Cold-anesthetized female *Ae. aegypti* (3 to 5 day-old) were intrathoracically inoculated with 69 nl of RPMI containing 100 particle forming unit (PFU) of WNV using a Nanoject II (Drummond Scientific Company) and needles made from 1.14 mm O.D. glass capillaries (1.75″ length, Drummond). Control mosquitoes were injected with the same volume of RPMI. After inoculation, mosquitoes were maintained under standard rearing conditions for 9 days before biting.

### Biting of mice

Mouse belly was shaved with a hair trimmer (VITIVA MINI, BIOSEB) one day before challenge. On the day of mosquito bite challenge, mice were anesthetized by intraperitoneal (IP) injection of 0.2 ml/mouse of ketamine (10 mg/ml, Imalgène 1000, Boehringer Ingelheim Animal Health) and xylazine (1 mg/ml, Rompun 2%, Elanco GmbH). A tape perforated with 2 mm openings was applied on trimmed skin to control mosquito bite location. 16h-starved mosquitoes were placed in small 3D-printed cages (Fig. S2) covered with mosquito net to allow biting. Mice were exposed to one to five encaged WNV-infected mosquitoes. Mosquito blood feeding was visually recorded and confirmed by the presence of blood in this abdomen. Only blood-fed mosquitoes were considered to confirm biting. Bite sites identified thanks to the openings in tape were marked with a permanent pen (Fig. S2).

### Tissue collection

2 mm skin biopsies were collected from the bite sites at 10 min, 1 h, 3 h, 6 h, 9 h, and 24 h post-bite using biopsy punches (Kai Medical) on mice anesthetized by IP injection of 0.2 ml/mouse of ketamine and xylazine. On the same mice, inguinal lymph nodes (LNs) and blood were collected.

### Absolute quantification of WNV (+) gRNA

Total RNA was extracted from skin biopsies, LNs, mosquitoes, SG or cells using EZNA RNA extraction kit I (OMEGA) or from blood samples using QIAamp Viral RNA Mini Kit (Qiagen). Absolute quantification of positive strand gRNA was performed through one-step RT-qPCR. Total reaction volume was 10 µl and contained 5 µl of iTaq Universal SYBR green one-step kit (Bio-Rad), 300 nM of forward and reverse primers ^44^ (Table S5) and 2 μl of RNA extract. The reaction was performed in AriaMx Real-time PCR System (Agilent) with the following thermal profile: 50°C for 10 min, 95°C for 1 min and 40 cycles of 95°C for 10 sec and 60°C for 25 sec, followed by a melting curve analysis. An absolute standard curve for DENV and WNV gRNA was generated by amplifying the qPCR target using primers detailed in Table S5 as previously performed ^143,144^. Using the gRNA templates, the limit of detection (LoD) at 95% was determined by calculating fractions of detected samples in six replicates of serial dilutions ^145^.

### Mouse symptom evaluation

After mosquito biting, mice were weighed daily to calculate the percent of weight loss. Daily clinical examinations were conducted and a clinical score (CS) ranging from 0 to 5 was assigned to each mouse following criteria from a previous study ^146^ where CS of 0 was assigned to healthy mice; CS of 1 for mice with ruffled fur, lethargy, hunched posture, no paresis, normal gait; CS of 2 for mice with altered gait, limited movement in 1 hind limb; CS of 3 for lack of movement, paralysis in 1 or both hind limbs, and CS of 4 for moribund mice. A CS of 5 indicated mortality (Table S1). Mice were euthanized if they displayed neurological symptoms, severe distress, or weight loss exceeding 20%. At day 15, mice were euthanized under anesthesia.

### Skin cell dissociation

Skin biopsies were minced with a scissor and incubated at 37°C for 60 min in 1 ml dissociation buffer containing 5 ml RPMI 1640 without Ca^2+^ or Mg^2+^ (No FBS or EDTA), 23.2 µl DNAse 1 (10U/µL) (Sigma-Aldrich), 50 µl Liberase (25mg/mL) (Sigma-Aldrich), 116 µl of 1M HEPES (Gibco), 116 µl of 100 mM sodium pyruvate (Gibco), 0.0025 g Hyaluronidase (Worthington), 500 µl of Dispase: Collagenase (1mg/ml) (Sigma-Aldrich). The dissociation was stopped by adding 10 µl of 0.5 M EDTA (Invitrogen) and 400 µl of FBS. Cell aggregates were mechanically dissociated by (i) vortexing for 30 sec, (ii) passing through 19G needle (TERUMO) several times (∼10 times), or (iii) using GentleMACS (MACS) dissociator with m_imptumor_01 program. Cells were filtered through 70 µm (PluriSelect) and 40 µm cell strainers before centrifugation at 500 g for 10 min at 4°C (low break speed) and resuspension in appropriate volume of PBS for flow cytometry or 0.04% BSA-RPMI for scRNA-seq.

### Flow cytometry

Three pools of 25 2-mm skin biopsies were collected from the trimmed abdomen of unbitten mice. After cell dissociation, cell surface staining of skin cells was performed in PBS buffer containing 1% FCS for 30 min at 4°C with the following antibodies: MHC Class II (I-A/I-E), monoclonal Antibody APC-eFluor780 (eBioscience), PE/Cyanine7 anti-mouse CD103 Antibody (Biolegend), APC anti-mouse/human CD207 (Langerin) Antibody (Biolegend). Cells were subsequently washed with PBS, pelleted by centrifugation at 500 x g for 5 min at 4°C, resuspended in PBS and analyzed on an LSR Fortessa (BD Biosciences) flow cytometer. Results were analyzed with FlowJo (BD Biosciences). Antigen-presenting cells (MHC-II+) present in skin cell suspensions were analyzed for CD103 and CD207 expression to evaluate the proportion of Langerhans cells (CD207+) and dermal dendritic cells (CD103+).

## 5′-end single cell RNA-sequencing

2-mm skin biopsies corresponding to the areas bitten by WNV-infected mosquitoes were collected at 6 h post biting. Each mouse was bitten by three to five mosquitoes. Twenty-two biopsies collected from eight animals were pooled. The same number of skin biopsies were collected from unbitten mice that underwent the same procedures as control (resting skin). Skin biopsies were cell dissociated using enzymatic digestion and needle-assisted mechanical dissociation and pelleted in 0.04% UltraPure BSA solution (Thermofisher Scientific). Dead cells were removed using the Dead Cell Removal Kit (Miltenyi Biotec) with MS columns. Live cells were resuspended in 2% UltraPure BSA, counted using Luna automated cell counter (Logos), and concentrated to ∼1,000 cell/µl. GEM generation, barcoding, post GEM-RT cleanup, cDNA amplification, and cDNA library construction were performed using the Chromium Single-Cell 5′ Reagent version 2 kit (10x Genomics). Libraries were sequenced on an Illumina HiSeq4000 platform (Illumina). Mouse challenge, cell preparation, GEM generation were performed at the BSL3 Vectopole of MIVEGEC, IRD, Montpellier; barcoding, post GEM-RT cleanup, cDNA amplification, library preparation in molecular biology laboratories at MIVEGEC; quality control and sequencing at the Genomics Research and Technology Hub of UCI. Multiple repeats were conducted and two repeats for infectious bite and one for resting skin were successful.

### Data processing and quality control for 5′-end transcripts

Raw counts were aligned against the *Mus musculus* (mm39), *Ae. aegypti* (AaegL5.0) ^147^, and West Nile Virus ^148^ genomes using 10X Genomics Cell Ranger v7.1.0. Data was preprocessed and integrated with Seurat ^149^. First, cells were removed if total RNA count was low (nCount_RNA < 500), number of expressed features was too low or too high (nFeature_RNA < 500 and nFeature_RNA > 6000), and mitochondrial gene count was high (percent.mt > 10). Second, the samples were normalized, clustered, and projected onto low-dimensional embedding (UMAP) following the standard Seurat workflow with default parameters. The first 20 PCA dimensions were used for computation of neighbors and low-dimensional embedding. Third, the samples (1 resting, 2 infectious bite) were integrated by selecting integration features (*SelectIntegrationFeatures()*) and anchors (*FindIntegrationAnchors()*), resulting in the integrated dataset (*IntegrateData()*). Finally, the integrated data was rescaled and UMAP representation was computed again. The two infectious bite samples were merged into a single sample for downstream analysis.

### Clustering, annotation, and proportion testing of 5’end transcripts

Cell types were identified by computing cell neighbors using the top 15 PCA dimensions, followed by clustering with resolution = 0.02, resulting in n = 8 clusters. This resolution threshold was identified by trial-and-error and based on the clear separation between the cell types in the UMAP space. Clusters were annotated based on the expression of marker genes identified with the *FindAllMarkers()* Seurat function with parameters (only.pos = TRUE, min.pct = 0.8, logfc.threshold = 0.5) based on Wilcoxon rank sum test and adjusted p-values with Bonferroni correction. Based on annotation at cell type level, fibroblasts, keratinocytes, lymphoid cells and myeloid cells were selected for individual sub-clustering. For each cell type, multiple clustering resolution parameters were tested to identify cell subtypes, and the final parameter value was set based on the interpretability of the resulting clusters (fibroblasts: 0.2; lymphoid: 0.8; myeloid: 0.9; keratinocytes: 0.3), whereby gene markers for each subtype were identified following the same procedure explained above. Proportion tests for cell types and subtypes between Resting and Infectious bite were performed using the scProportionTest R package ^150^ by applying a permutation test that associates a p-value to the cell fraction fold-change ^150^.

### Identification of mosquito RNA-containing cells

Based on alignment against the *Ae. aegypti* genome (AaegL5.0) ^147^, cells were categorized as mosquito-positive or mosquito-negative based on detection of one or more mosquito RNA reads, thus enabling the quantification of mosquito-positive cells per cell type and per subtype (in the case of fibroblasts). Further classification based on the number of mosquito RNA reads was not conducted as the majority of cells expressed either none or a single mosquito RNA reads.

### Identification of differentially expressed genes (DEGs)

DEGs between mosquito RNA-positive and -negative cells within fibroblasts, and between resting and infectious samples within cell subtypes were identified with the *findmarkers()* function from Seurat. Given the sparsity of several genes with high fold-change in the infectious bite, we implemented the MAST statistical testing that specifically considers zero-inflation in scRNA-seq data ^151^ by setting *test.use=”MAST”*. This statistical test was performed for all main cell types as well as on the subtypes for fibroblasts, keratinocytes, lymphoid cells and myeloid cells. DEGs were defined based on adjusted p value < 0.001 and a |log2 fold-change| > 2. DEGs were functionally categorized into immunity (including inflammation, innate immunity, adaptive immunity), skin regeneration (including cell motility, cell proliferation, angiogenesis, tissue development, ECM), metabolism, RTT (replication, transcription and translation), Redox–stress–mitochondria, non-coding RNA, or divers (for genes not related to the above functions or with unknown functions) based on GO biological processes retrieved from Ensembl database (ensembl.org/biomart/martview/).

### Gene Ontology (GO) enrichment analysis

GO enrichment analysis was performed on the within cell subtypes upregulated DEGs with the gseapy package ^152^ using the ‘GO_Biological_Process_2025’ gene sets. Significant biological processes were filtered based on GO term presence in > 10% of within-subtype DEGs and p-value < 0.01 (i.e., log10 (1/FDR) > 2). Combined score represents the product of the p-value and the z-score of the deviation from expected rank, providing a comprehensive measure of enrichment strength.

### Cell-cell communication analyses

The data was analyzed using the CellChat package^153^. First, the individual samples (Rest, Infectious bite) were converted into CellChat data objects following the standard procedure explained in CellChat tutorial. Cell-cell communication analysis was also performed using CellChat’s default setting, except for setting a minimum number of cells min.cells = 5 per cell type to filter out weak cell-cell communication pathways (i.e., the pathway must be detected in at least 5 cells per cell type). Then, the samples were integrated, and the cell-cell communication strength and pathways were identified and plotted using the netVisual_heatmap and Ranknet functions as well as custom script.

### Collection of mosquito saliva

Bulk saliva collection was based on previously described protocols ^154,155^. Uninfected female mosquitoes were let to feed on a Hemotek feeding system (Hemotek) containing 3 ml of RPMI and 1% 25 mM Erioglaucine (SigmaAldrich). The number of mosquitoes that salivated was estimated by visually counting individuals with blue abdomens, indicative of feeding and salivation. Salivation media was collected after 1h.

### Evaluation of cell viability

10^5^ HFF-1, U-937 or Jurkat were supplemented or not with 1% Erioglaucine. After 6h, cell viability was estimated using CyQUANT NF Cell Proliferation Assay Kit (Invitrogen).

### Cell infection

10^5^ HFF-1, U-937 or Jurkat cells were seeded in 48-well plates. 24h later, cells were treated with WNV at MOI = 1, with or without the addition of saliva collected from 5 mosquitoes (4.5 µl of solution corresponding to saliva from one mosquito), or with mosquito saliva alone. Control cells received the same volume of saliva collection medium. At 6 h post treatments, cells were collected in TRK lysis buffer.

### Relative gene and WNV gRNA quantification

Total RNA from cells was extracted using EZNA Total RNA kit I. Relative gene expression was quantified with one-step RT-qPCR using iTaq Universal SYBR green one-step kit (Bio-Rad) with the corresponding primers (Table S1). Amplification was conducted in AriaMx using the same thermal profile as for WNV (+) gRNA quantification. Expression of three housekeeping genes (i.e., *RPL13A, ACTB,* and *GAPDH*) was used for normalization (Table S5). Relative expression was calculated using the 2-ΔΔCt method.

### siRNA silencing

10^5^ U-937 or HFF-1 cells were transfected with 12 pmol of multiplex flexitube siRNAs (Qiagen) using Lipofectamine RNAiMax (ThermoFisherScientific). siRNA GS131578 against *Lrrc15* was used in HFF1 cells; siRNA GS10437 against *ifi30* and siRNA GS3123 against *Hla-drb1* were used in U-937 cells (Table S1). Allstars multiplex siRNAs (Qiagen) was transfected as control. 48 h after transfection, cells were infected with WNV as described above. At 36 hpi, cells were collected in TRK lysis buffer for RNA quantification or in RIPA lysis buffers (ThermoFisherScientific) protein analysis.

### siRNA delivery in skin

One day prior siRNA injection, mouse hairs on the lower back were shaved with the hair trimmer (VITIVA MINI, BIOSEB). Anesthetized mice were intradermally inoculated, using CellTram 4r Oil microinjector (Calibre Scientific), with 30 pmol in 3 µl of siRNA GS74488 against *Lrrc15* or Allstars multiplex siRNAs (Qiagen) mixed with 5 µl of Lipofectamine RNAiMax, completed to 10 µl with water. Each mouse was injected at three different sites around the planned biting site. The inoculation sites were identified by a permanent pen.

### Western blot on cells and skin

48 h post siRNA transfection, cells and skin were collected. Cells were washed twice with PBS, scrapped in 70 μl of RIPA containing 1X protease inhibitors (Sigma-Aldrich) and the supernatant was collected after centrifugation at 12,000 g for 1 min. For each mouse, ∼5 mm diameter biopsies were collected from the three siRNA injected-bitten skin in 200 µl 1X RIPA lysis buffer, micro-dissected with scissors, homogenized using Fast-prep bead bitter (MP) with glass beads, and supernatant was collected after centrifugation at 12,000 g for 1 min. Normalized protein quantities were separated under denaturing conditions in 10% polyacrylamide gel and transferred onto 0.2 µm Nitrocellulose membrane using TransBlot system (BioRad). Staining was conducted with 1:1,000 and 1:500 rabbit anti-LRRC15 (50546, Cell Signaling) for human cells and mouse skin, respectively, and 1:400 anti-Actin (MA5-11869, Invitrogen) in 0.1 % Tween-20 1% BSA in PBS. Secondary staining with 1:2,000 of goat anti-rabbit (7074P2, Cell Signaling) and goat anti-mouse (7076P2, Cell Signaling) was similarly performed. Pictures were taken with ChemiDoc (BioRad) using SuperSignal West Pico Chemiluminescent substrate (ThermoFisherScientific). Actin expression was used for normalization.

### Quantification of siRNA silencing effect on skin infection and symptoms

48 h post siRNA injection, mice were bitten at the siRNA injection sites by one WNV-infected mosquito. Each mouse was bitten by three mosquitoes at three different sites previously injected with siRNA. At 24h post-bite, skin biopsies from the bite sites and inguinal LNs were collected and homogenized with Fast-prep bead bitter (MP) with glass beads in 350 µl of TRK lysis buffer (EZNA). After RNA extraction with EZNA Total RNA extraction kit I, WNV gRNA was quantified. Additional mice were monitored for symptoms, weight loss and survival. At 4 days post bite, blood samples were collected via mandibular puncture and sample volumes were estimated by pipetting before RNA extraction using EZNA total RNA extraction kit I and WNV gRNA quantification. Mice were euthanized if they displayed neurological symptoms, severe distress, or weight loss exceeding 20%. Mice were euthanized under anesthesia at day 14.

### Skin Human Cell Atlas (HCA) data meta-analysis

We re-analyzed the skin component of the HCA, which is an integrated single-cell dataset from 24 independent single-cell datasets of normal, non-diseased human skin sampled across different ages, sexes and genders, and anatomical body sites. Skin HCA contains two levels of cell annotations. The first level of annotations identifies all skin-resident cell lineages. The second level identifies lineage-specific subclusters. We analyzed *LRRC15* expression across all lineages, before restricting analysis to fibroblast subclusters. Before analyzing *LRRC15* expression in fibroblasts across anatomical regions, we removed cells sampled from anatomical regions for which there were fewer than 500 fibroblasts, and cells from anatomical regions of unknown origin. This step of filtering was to ensure that observations of low *LRRC15* expression was a true biological signal, rather than due to low fibroblast numbers.

### Spatial mapping of skin HCA to Xenium data

Prior to spatial projection analysis, we optimized SpaceRanger-generated cell segmentation of the raw spatial Xenium data using ProSeg ^156^. We then performed quality control of individual samples using Spotsweeper-py ^157^, removing cells with excessively low total counts, excessively low numbers of unique transcripts, and an excessively high proportion of gene transcription due to mitochondrial genes. We then normalized cell counts by median library size before log-transforming the normalized counts (adding a pseudocount of 1). When analyzing the Scalp sample, we used ResolVI ^158^ to cluster the data initially and remove adipocytes, as this cell lineage is absent from single-cell skin HCA dataset.

We used cell2location ^159^ to project HCA fibroblast populations to their likely spatial location in Xenium spatial transcriptomics samples. We assumed that each Xenium cell mapped exactly to one cell type. As cell2location was originally developed for the first generation 10X Visium spatial sequencing platform, which contains multicellular spots, cell2location output estimates the expected “number” of cells from each cell annotation label at each spot. Therefore, we annotated Xenium cells as the cell type annotation with the highest number of cells predicted by cell2location. Briefly, we performed two levels of spatial projection using the HCA and the Xenium samples. First, we trained cell2location using the global cell lineage annotations to predict which Xenium cells were likely fibroblasts. Then, we trained a second cell2location model using only HCA fibroblast subcluster annotations and projected these annotations onto the identified Xenium fibroblast cells.

### Statistical analysis

Differences in log-transformed gRNA copies and relative gene expression were tested with one-tailed T-test or post-hoc Dunnett’s test. Differences in survival were tested with Kaplan-Meier survival analysis with Gehan-Breslow-Wilcoxon test. The statistical analyses were performed using Prism 8.0.2 (GraphPad). Clinical scores and weight loss were analyzed using generalized linear mixed-effects models (GLMMs). Clinical scores were modeled assuming a Poisson error distribution with a log link function. Weight loss were modeled using an inverse Gaussian error distribution with a log link. Both models included dpi, condition (siNeg vs. siLRRC15), and their interaction as fixed effects, with mouse specified as a random intercept. Analyses were performed in R (version 4.3.3) using the glmmTMB package (version 1.1.9). Model adequacy was evaluated using simulation-based diagnostics implemented in the DHARMa package (version 0.4.7). No evidence of overdispersion was detected for either the clinical score model (dispersion = 0.579, p = 0.496) or the weight loss model (dispersion = 1.038, p = 0.728).

## Data and Code Availability

scRNA-seq data have been deposited at https://zenodo.org/records/17953412 and are publicly available as of the date of publication. Any additional information required to reanalyze the data reported in this paper is available from the Lead Contact upon request.

## Acknowledgments

We thank all members of the Pompon’s Laboratory at MIVEGEC, IRD, for their valuable suggestions and support. We are especially grateful to VectoPole team in Montpellier, particularly Bethsabée Scheid and Carole Ginibre, for providing mosquito eggs, and to Nathalie Barougier, Pascal Bautinaud and Drs. Sylvie Cornelie and Idris Mhaidi for their assistance with mouse management. We thank Dany Sevrac and the MGX-Montpellier GenomiX platfrom, part of the French National Infrastructure at the Montpellier Biocampus, for providing the Chromium system and technical support. We acknowledge the imaging facility MRI, member of the national infrastructure France-BioImaging (https://ror.org/01y7vt929) supported by the French National Research Agency (ANR-24-INBS-0005 FBI BIOGEN). Finally, we are grateful to Melanie Oakes and the Genomics Research and Technology Hub at the University of California, Irvine (UCI), USA, for sequencing support.

