## Supplemental material for "Single-cell characterization of skin response to a bite by West Nile virus-infected mosquito reveals fibroblast-mediated barrier to transmission"

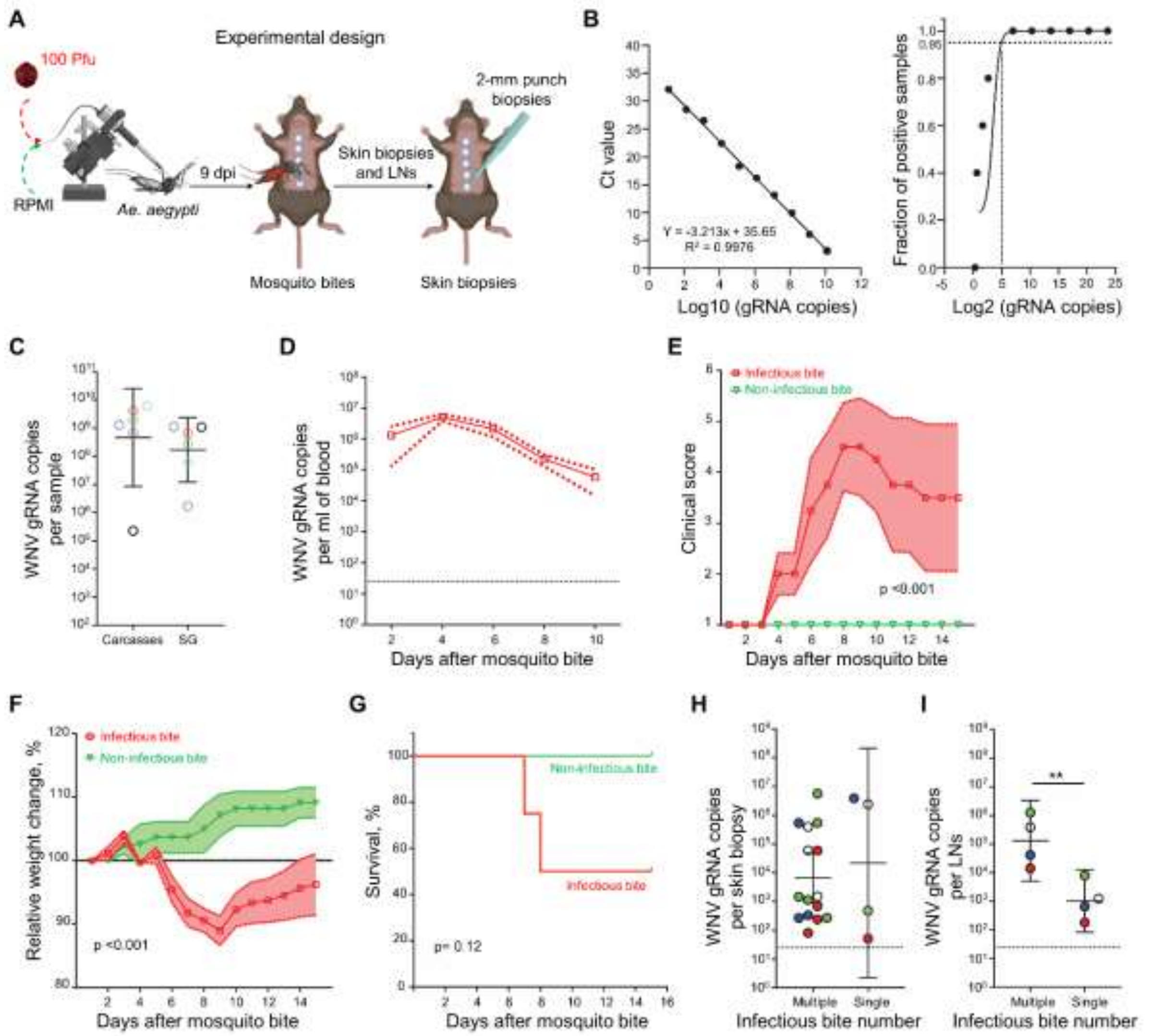

**Figure S1. Characterization of the mouse transmission model bitten by WNV-infected mosquitoes, related to Figure 1**

**(A)** Schematic depiction of the WNV-infected bite-initiated transmission mouse model. **(B)** Absolute quantification of WNV gRNA. Left graph, standard curve for gRNA absolute quantification. Points represent the averaged Ct values calculated from 4 repeats. Right graph, Limit of Detection (LoD) at 95% confidence for WNV gRNA is 25 copies ($5^2$). Fractions of positive samples were computed based on 4 or 5 replicates. **(C)** WNV gRNA copies in carcasses and salivary glands (SG) of the same mosquito at nine days after WNV injection. Corresponding mosquito samples are color-coded. **(D-G)** Mean viremia (D), mean clinical scores (E), mean relative weight changes (F), and survival (G) over 15 days after infectious (WNV) vs. non-infectious bite.

**(H, I)** WNV gRNA copies per skin biopsy (H) and pair of inguinal LNs (I) at 24 hours after either multiple or a single infectious bite. Points indicate different skin and LN repeats from 4 mice. Corresponding samples from the same mouse are color-coded. (C, H, I) Geometric means ± 95% CI. $**$, $p < 0.01$ by T test. (D, E, F) Mean ± s.e.m. (E, F) p indicates statistical differences for time × infection interaction. (G) p shows Kaplan-Meier survival analysis.

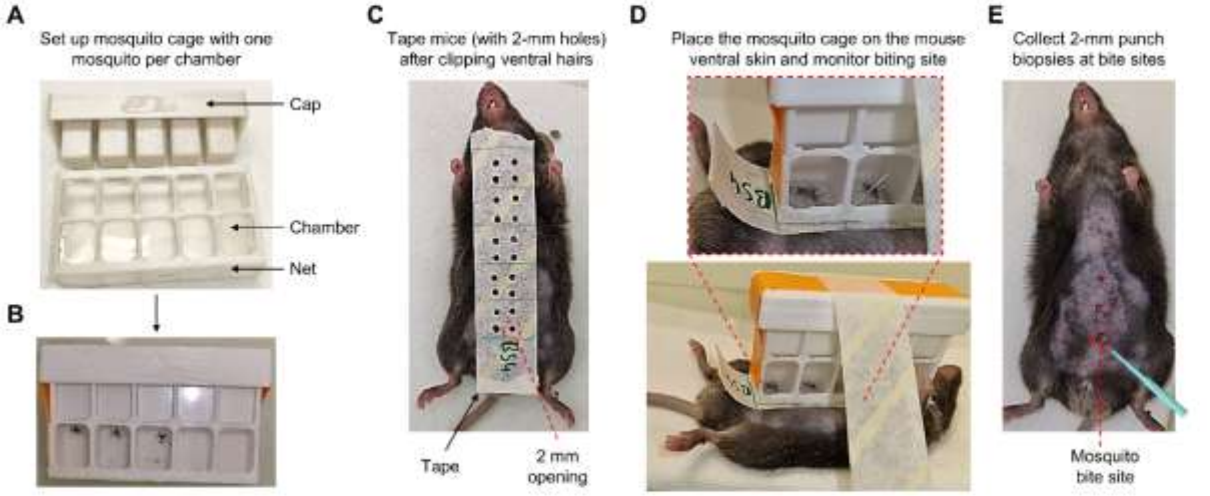

**Figure S2. Experimental design for identification of mosquito bites, related to Figure 1**

**(A)** An example of 3D-printed cage covered with mosquito net to allow controlled biting access.

**(B)** An example of starved mosquitoes placed in the cage, which is then sealed with the tape.

**(C)** An example of perforated tape with 2 mm openings applied onto abdominal skin of anesthetized mouse to direct mosquito bite location.

**(D)** Demonstration of the mosquito cage being secured onto mouse abdomen with mosquito biting activity being monitored visually.

**(E)** Demonstration of how individual bite sites were being identified and marked through the tape openings before biopsy collection.

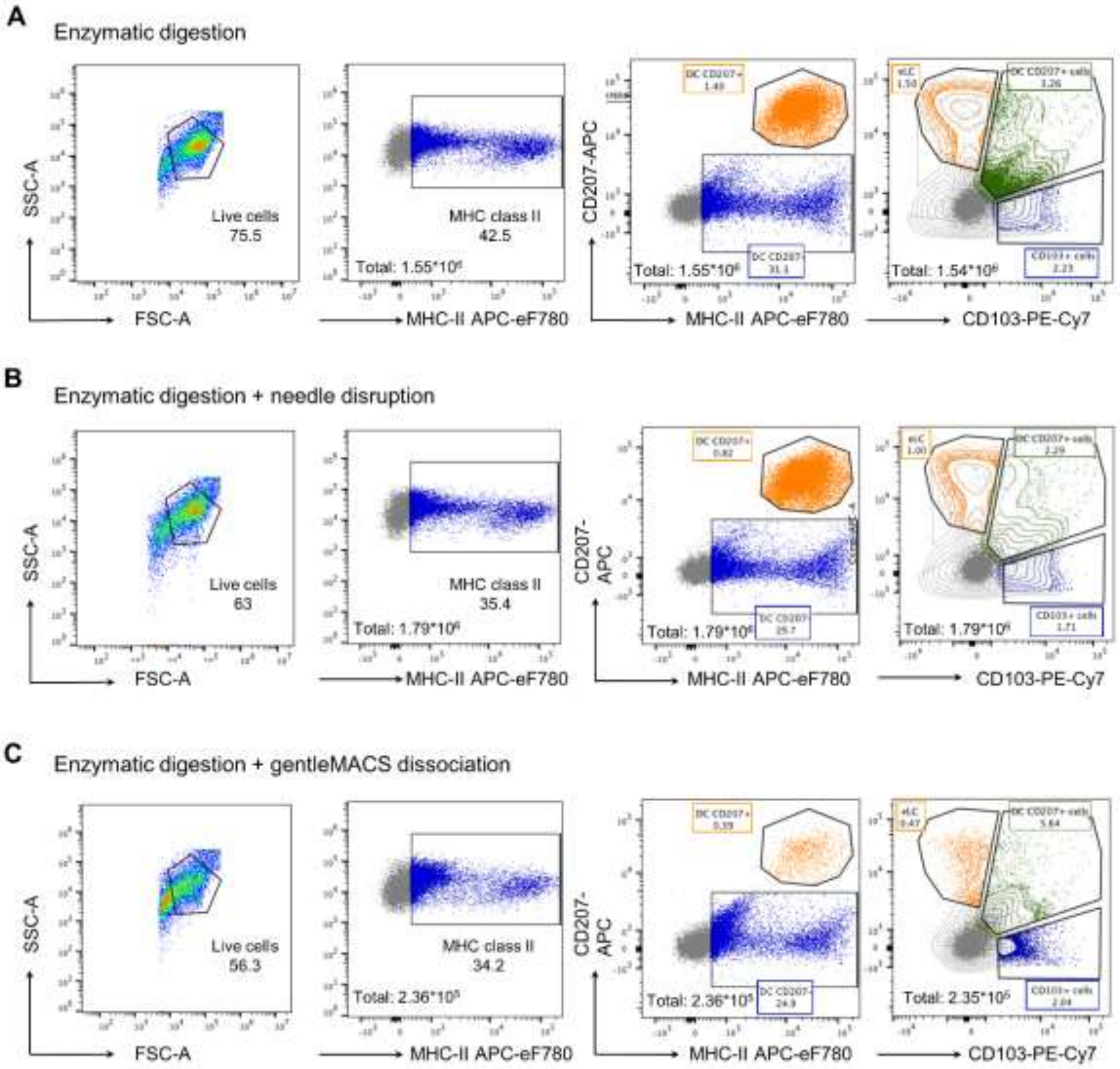

**Figure S3. Enzymatic skin tissue digestion combined with syringe disruption yields the highest number of cells, related to Figure 1**

**(A-C)** Quantification of DCs from the skin after enzymatic digestion alone (A), enzymatic digestion followed by needle disruption (B) and enzymatic digestion combined with gentleMACS-based dissociation (C).

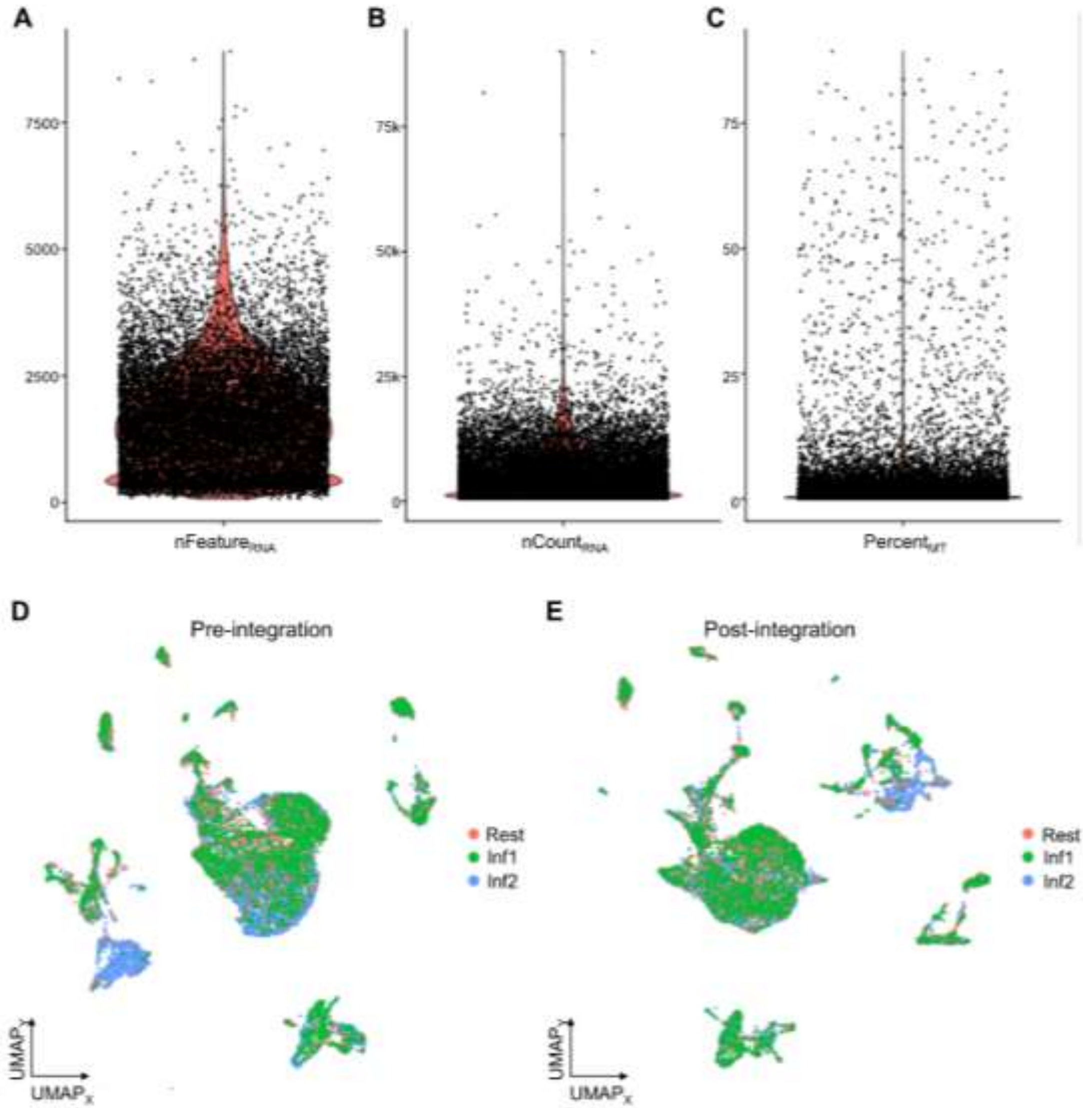

**Figure S4. scRNA-seq data pre-processing, related to Figure 1**

**(A-C)** Quality control metrics including number of RNA features (A), total RNA counts (B), and percent of mitochondrial counts (C) per cell for all samples.

**(D, E)** Samples (1 resting, 2 infectious bites) in low-dimensional UMAP embedding before and after integration.

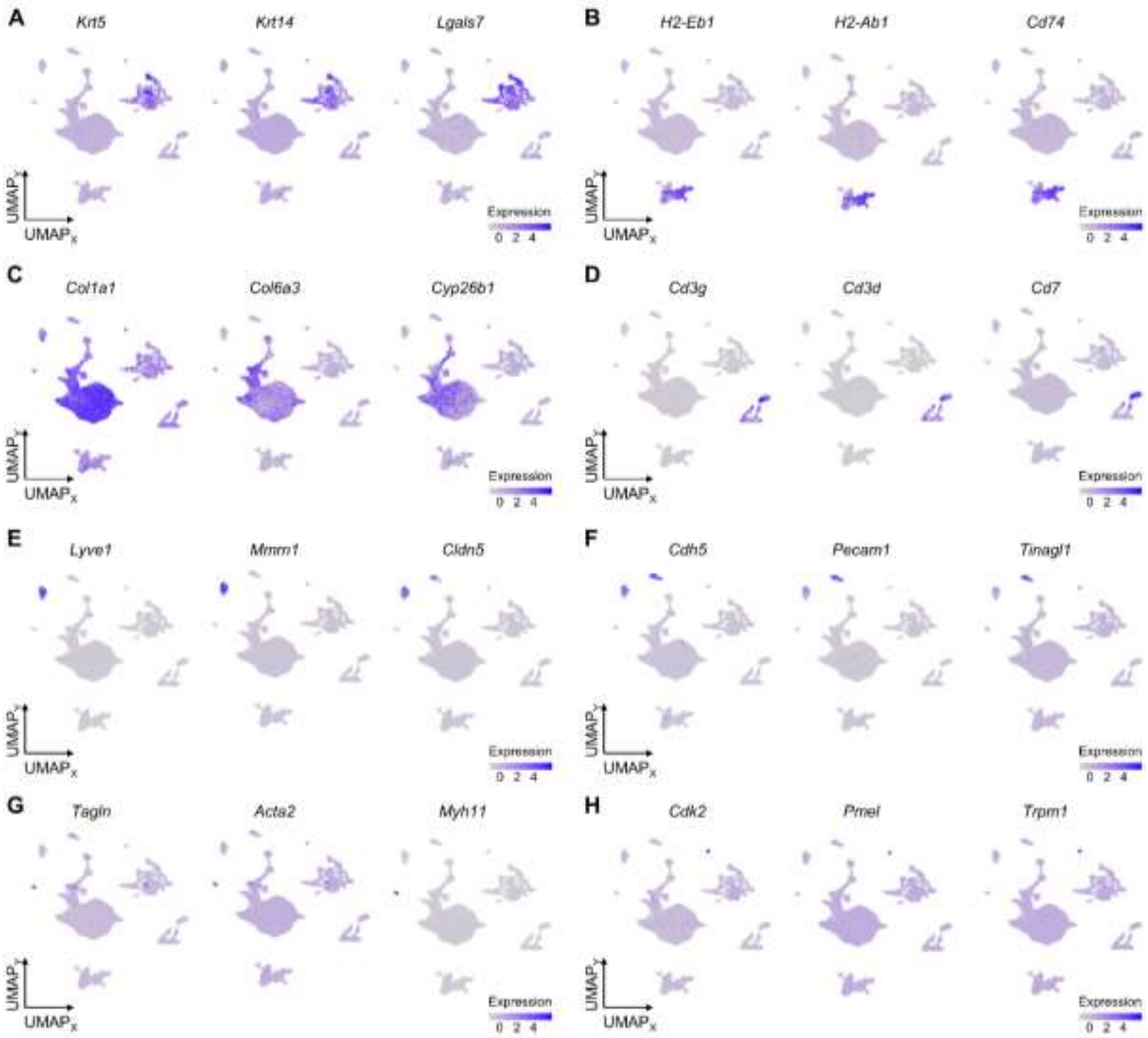

**Figure S5. Feature plots of key marker genes used for cell type annotation, related to Figure 1**

(**A-H**) Low dimensional embedding feature plots of key cell marker genes for keratinocytes (A), myeloid cells (B), fibroblasts (C), lymphocytes (D), lymphatic endothelial cells (E), vascular endothelial cells (F), vascular smooth muscle cells (G) and melanocytes (H).

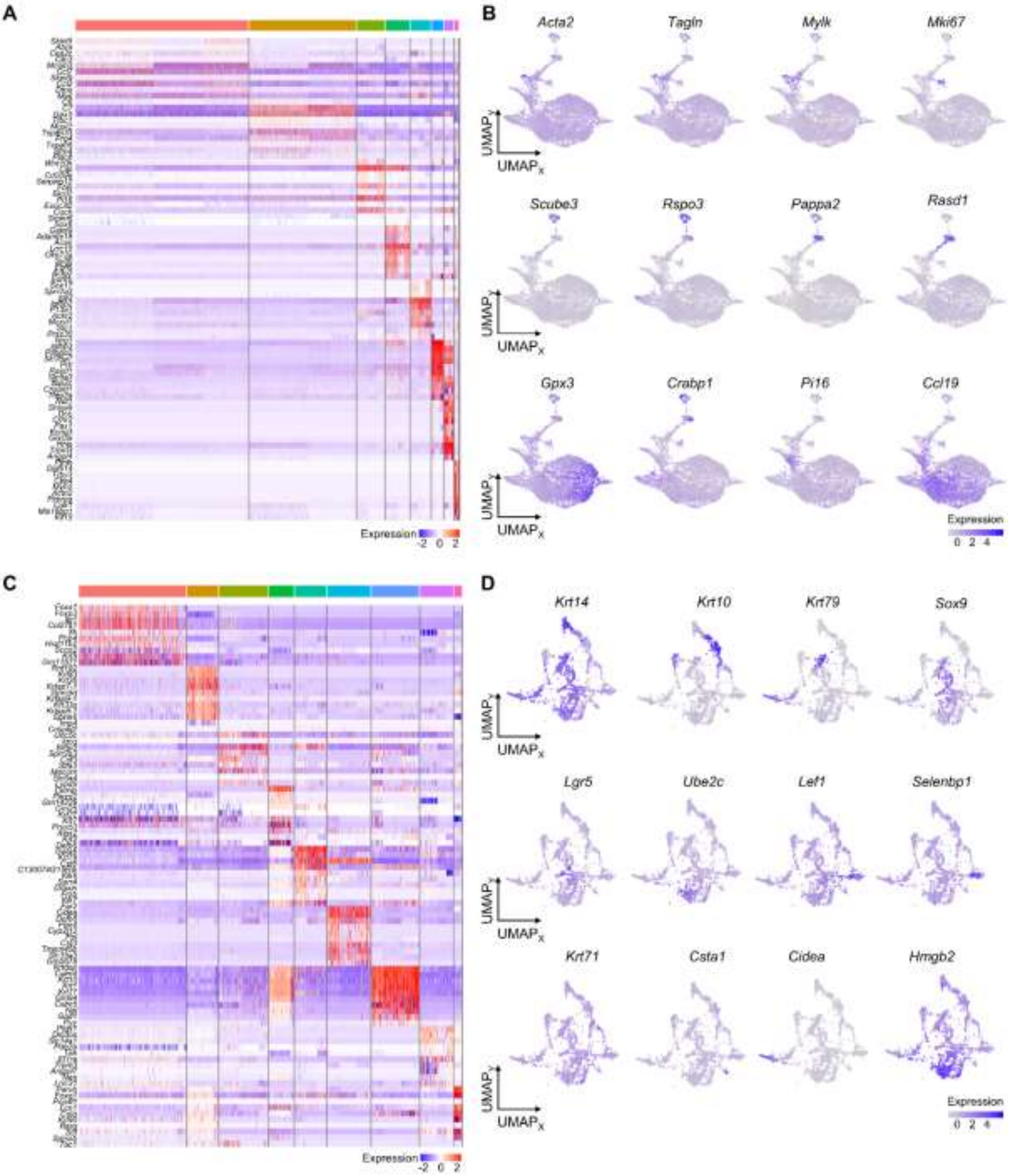
A
B
Acta2
Tagln
Mylk
Mki67
Scube3
Rspo3
Pappa2
Rasd1
Gpx3
Crabp1
Pi16
Ccl19
UMAPY
UMAPX
Expression
-2 0 2
0 2 4
C
D
Krt14
Krt10
Krt79
Sox9
Lgr5
Ube2c
Lef1
Selenbp1
Krt71
Csta1
Cidea
Hmgb2

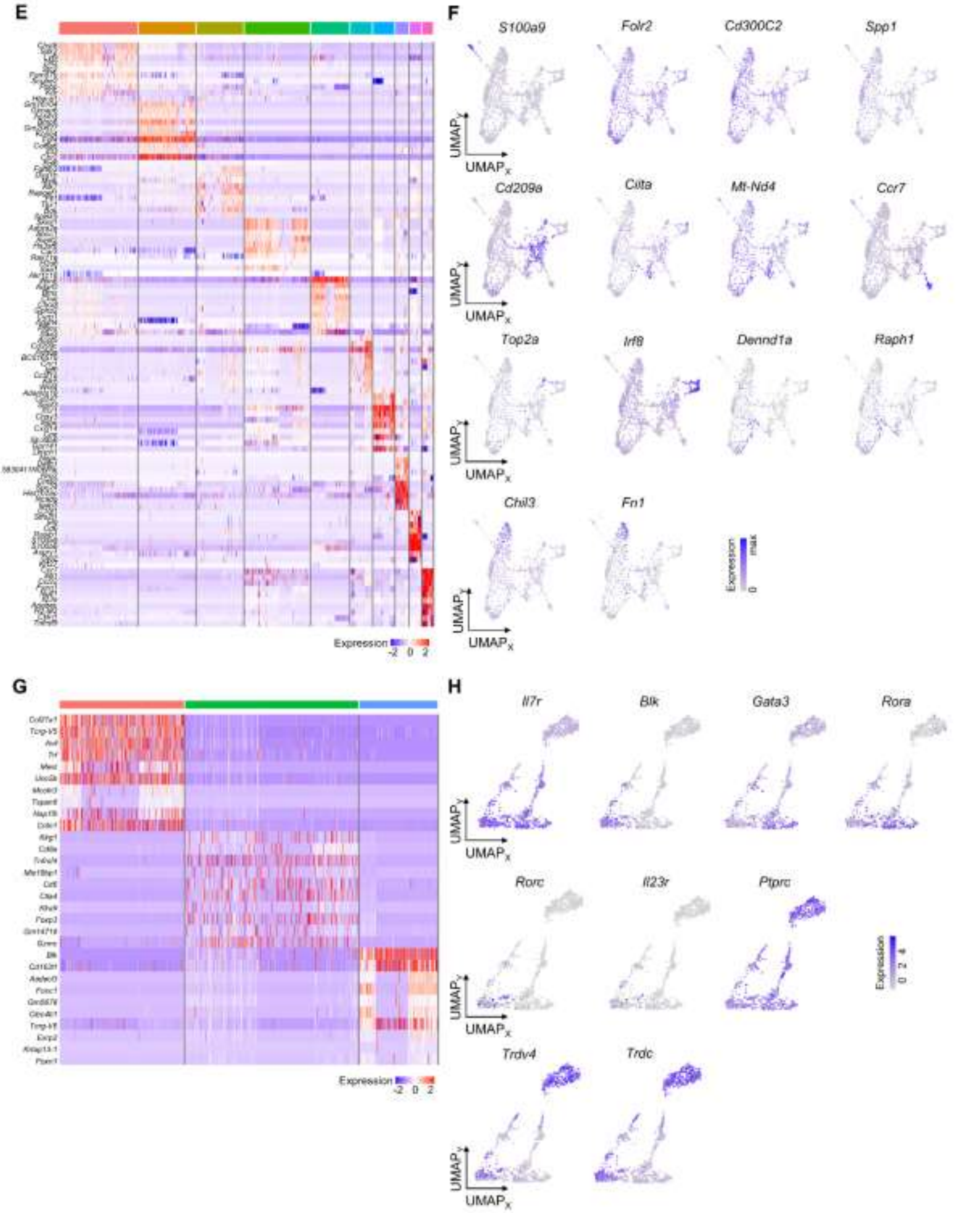

**Figure S6. Analysis of subcluster-specific marker genes, related to Figure 1**

**(A-H)** Heatmaps and feature plots of marker genes for cell subtypes obtained upon subclustering of fibroblasts (A, B), keratinocytes (C, D), myeloid cells (E, F), and lymphocytes (G, H).

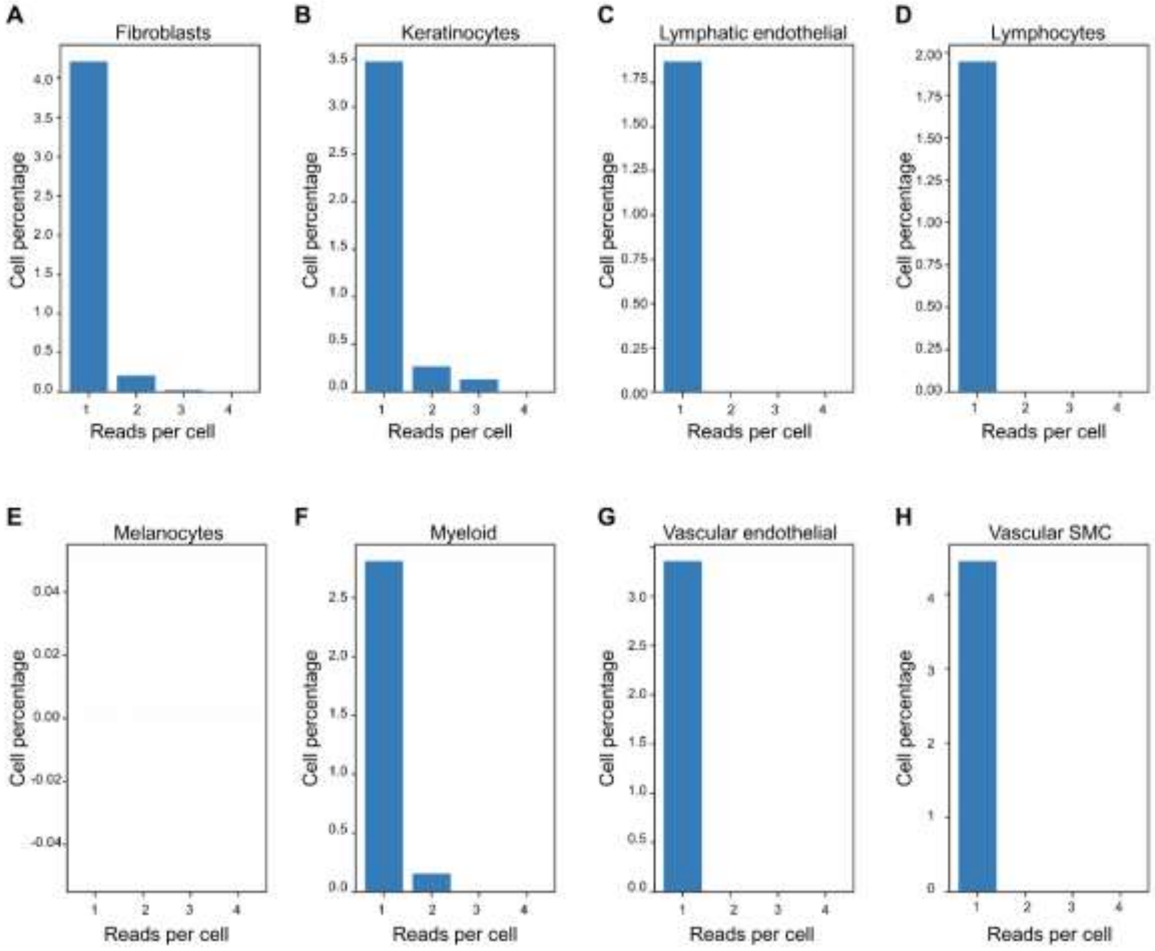

**Figure S7. Quantity of mosquito RNA reads per mouse cell type, related to Figure 2**

**(A-H)** Percent of cells with 1-to-4 mosquito RNA reads among fibroblasts (A), keratinocytes (B), lymphatic endothelial cells (C), lymphocytes (D), melanocytes (E), myeloid cells (F), vascular endothelial cells (G) and vascular smooth muscle cells (H). None of the cells had more than 3 RNA reads.

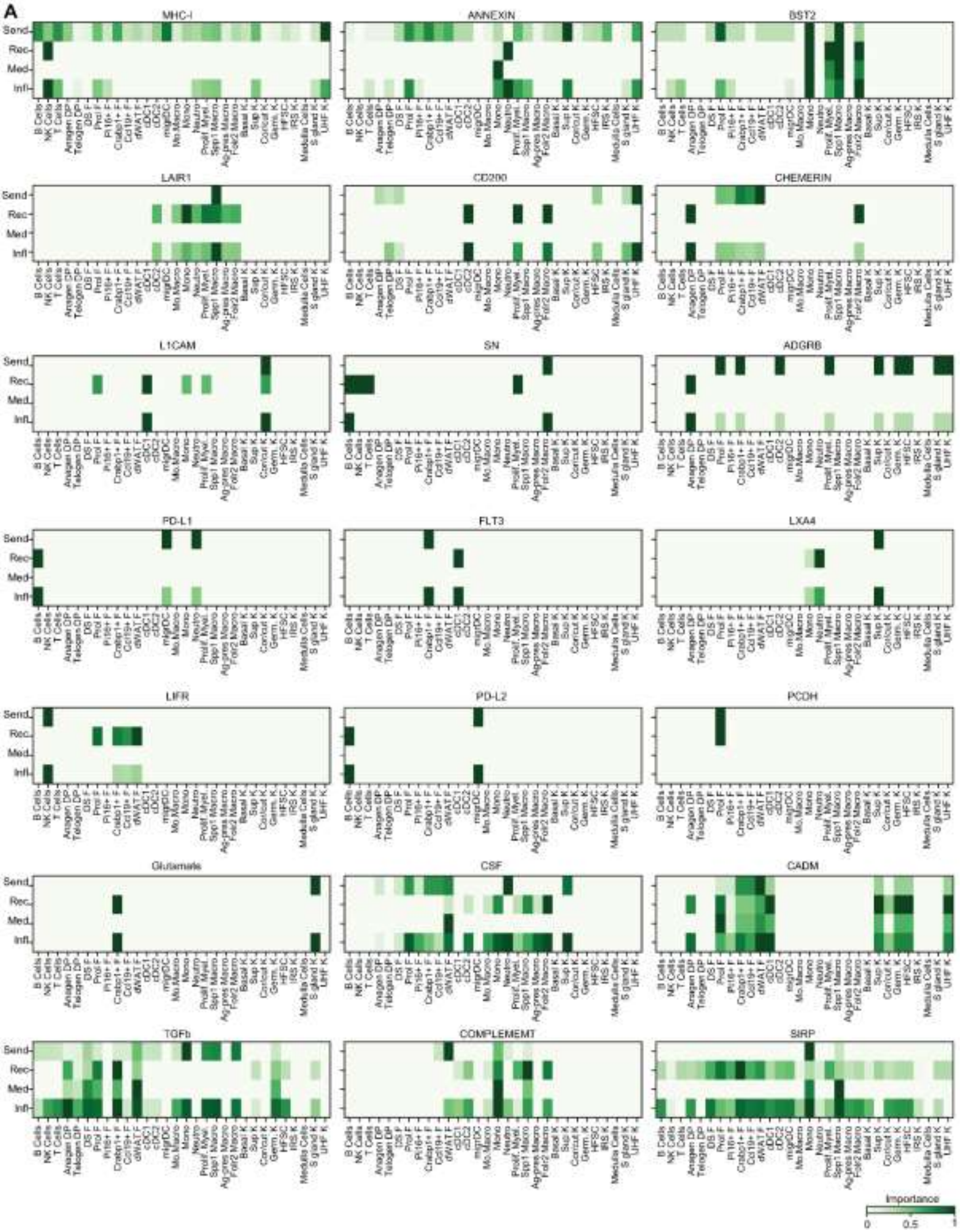
A
MHC-I
ANNEXIN
BST2
LAIR1
CD200
CHEMERIN
L1CAM
SN
ADGRB
PD-L1
FLT3
LXA4
LIFR
PD-L2
PCDH
Glutamate
CSF
CADM
TGFb
COMPLEMEMT
SIRP
Send
Rec
Med
Infl
Importance
0
0.5
1

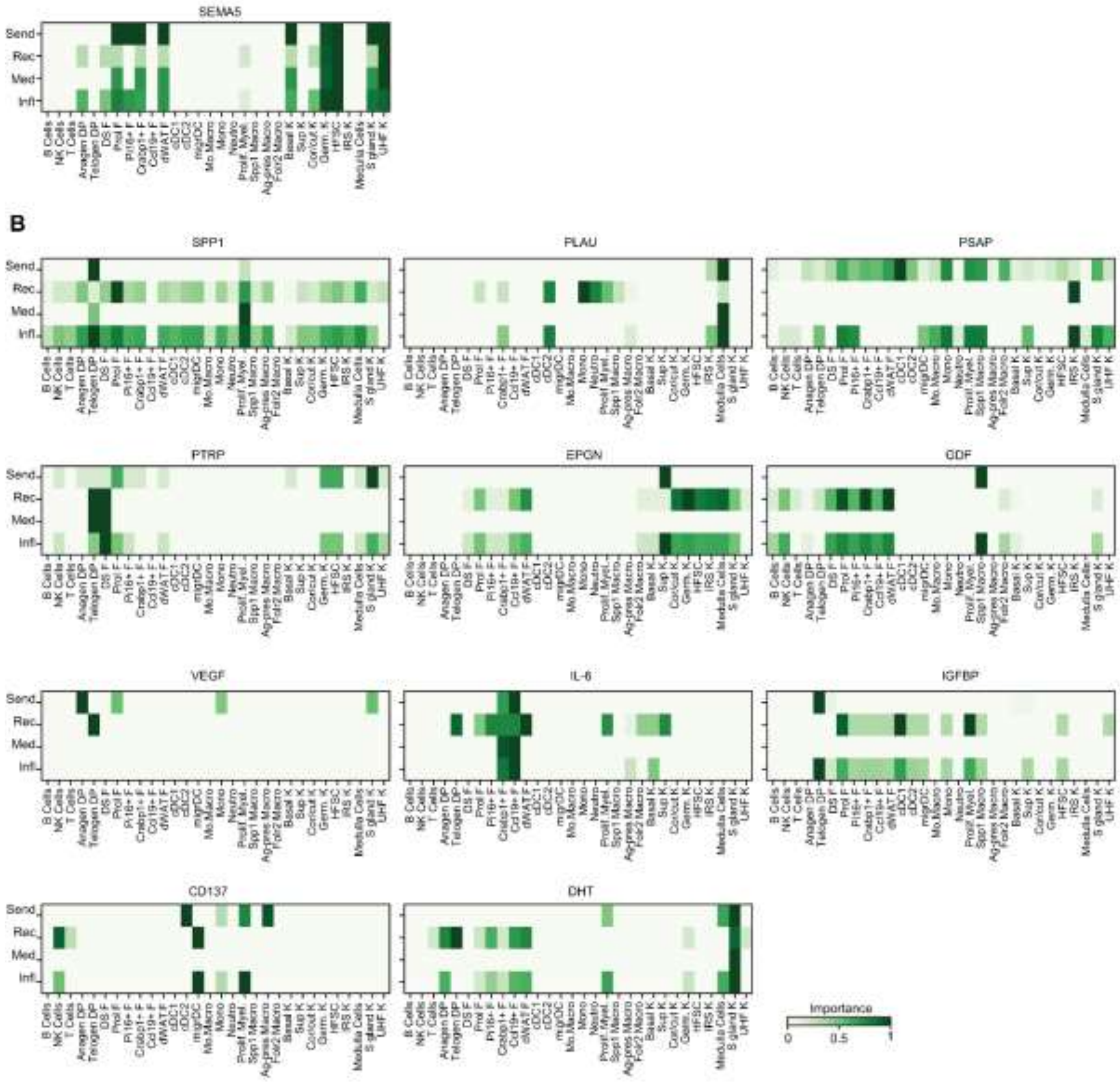

**Figure S8. CellChat analysis on cell-cell communication, related to Figure 4**

**(A, B)** Signaling roles for each cell subtype for the 22 upregulated (A) and the 11 downregulated (B) cell-cell communication pathways in response to infectious bite. NK, natural killer; F, fibroblasts; DP, dermal papilla; DS, dermal sheath; migr, migratory; mo.Macro, monocyte-derived macrophages; Mono, monocytes; Prolif., proliferative; Macro, macrophage; Neutro, neutrophil; K, keratinocytes; Sup, suprabasal; Cor/cut, cortex/cuticle; Germ, germinative; HFSC, hair follicle stem cells; IRS, inner root sheath; S, sebaceous; UHF, upper hair follicle.

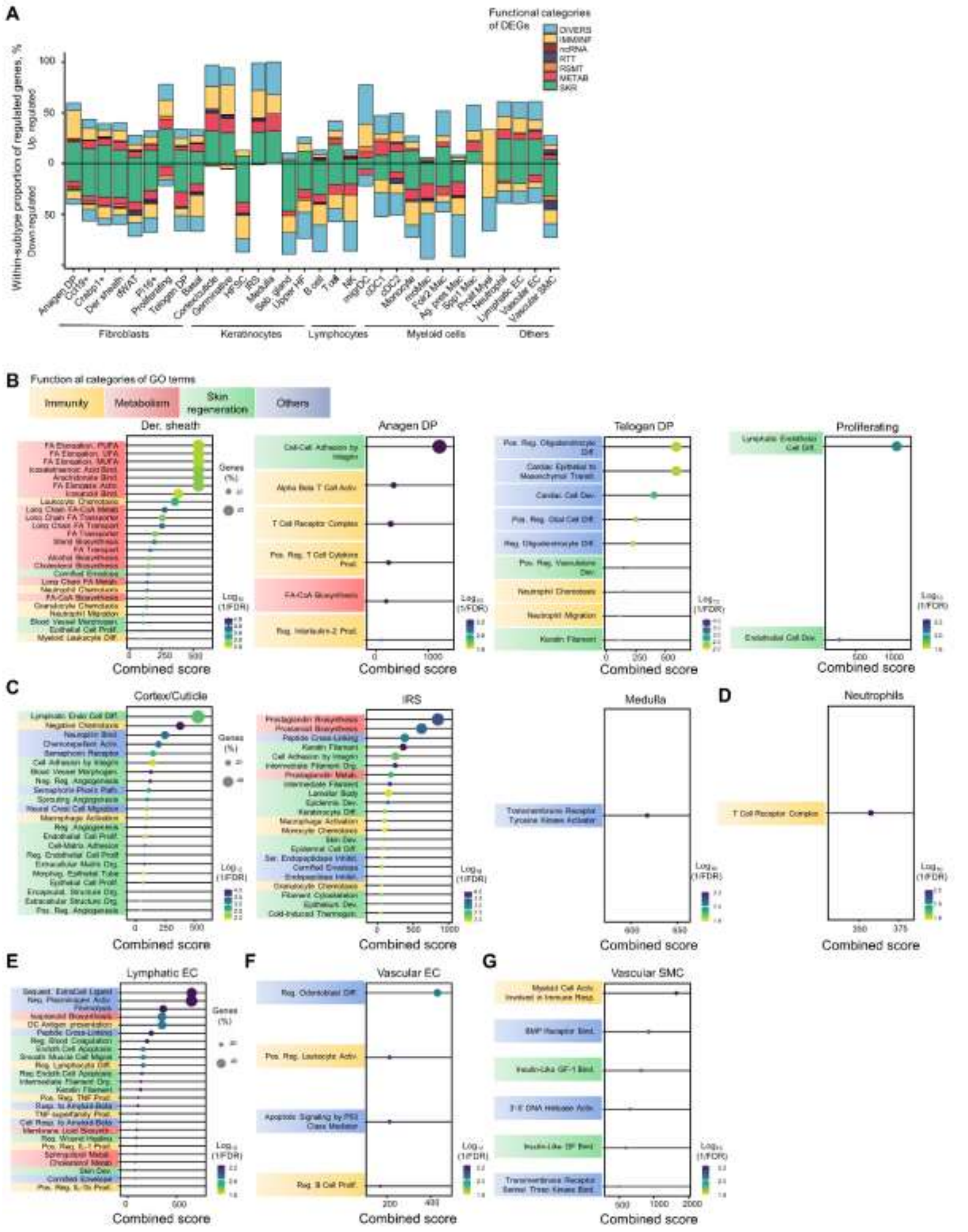

**Figure S9. Functional characterization of cell-specific transcriptomic responses to infectious bite, related to Figure 5**

**(A)** Functional distribution of differentially expressed genes (DEG) (adj. p-value < 0.001; |fold change| ≥ 4) within cell subtypes. Bars indicate percent of regulated DEG within cell subtypes. Functional categories include: SKR, skin regeneration; METAB, metabolism; RSMT, redox-stress-mitochondria; RTT, replication, transcription and translation; ncRNA, non-coding RNA; IMM/INF, immunity and inflammation; DIVERS, other and unknown functions.

**(B-F)** Gene ontology (GO) biological process enrichment analysis for the remaining subtypes of fibroblasts (B), keratinocytes (C), lymphatic endothelial cells (D), vascular endothelial cells (E) and vascular smooth muscle cells (F). Cell subtypes and main cell types are presented in an increasing order of abundance. Significant GO terms encompass > 10% of all differentially expressed genes within cell subtypes and have p-value < 0.01. Cell subtypes/main cell types that are not represented in either Fig. 5 or here did not have any significantly enriched GO terms. Functional categories are indicated by colored boxes. DP, dermal papilla; EC, endothelial cells; SMC, smooth muscle cells.

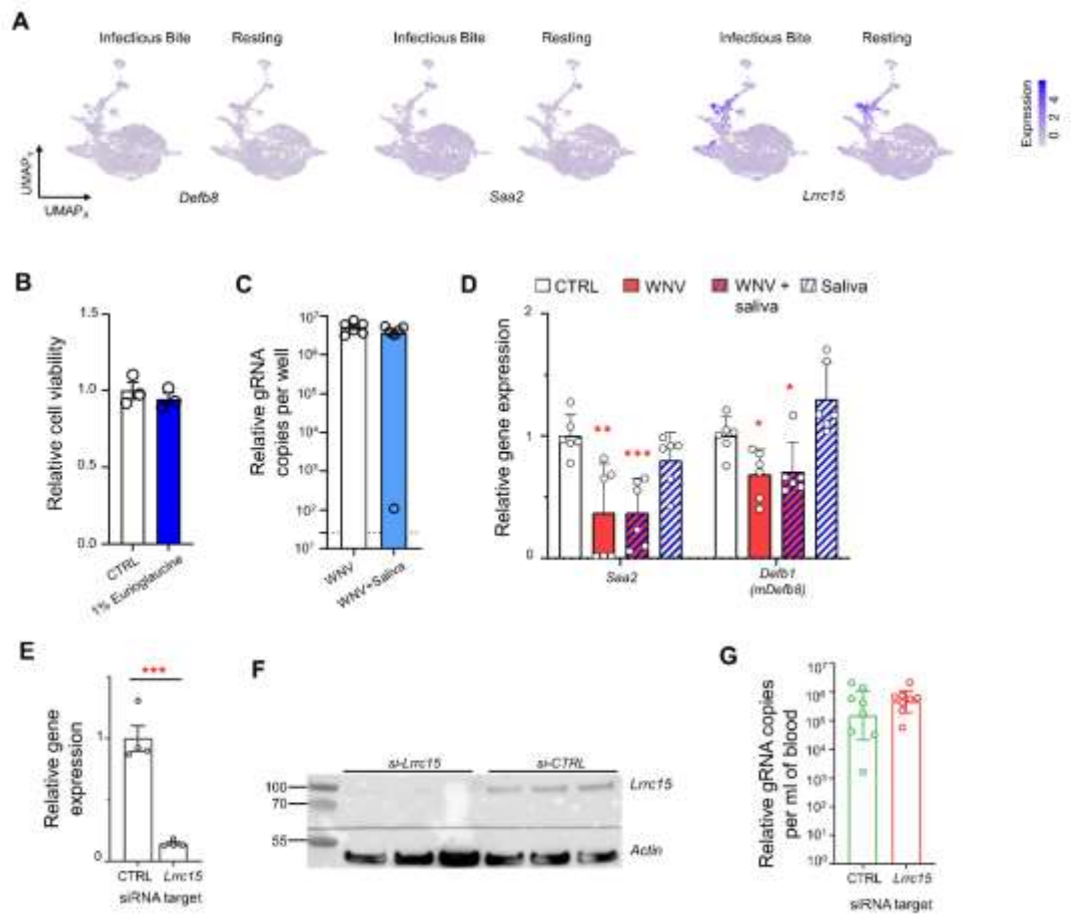

**Figure S10. Functional evaluation of skin cell responses, related to Figure 6**

**(A)** UMAP plots of the selected differentially expressed genes (DEG) after infectious bite vs. resting in fibroblasts.

**(B)** HFF1 cell viability at 6 hours post-treatment with 1% Erioglaucine. CTRL, untreated cells.

**(C)** Infection levels in HFF1 cells at 36 hours after infection with or without mosquito saliva supplementation.

**(D)** Regulation of expression of selected DEGs by either saliva, virus (WNV) or both in HFF1 cells at 6 hours post-treatment. CTRL, mock infection. These DEGs were differentially regulated in cell lines and skin.

**(E)** Expression levels of *Lrrc15* in si-Lrrc15-transfected HFF1 at 36 hours post-injection. CTRL, siRNA control.

**(F)** WB of Lrrc15 at 48 hours after siRNA transfection in si-Lrrc15-tranfected HFF1 cells. Actin level was used for normalization. CTRL, siRNA control. **(G)** RNAemia in mice at 4 days after mosquito challenge. N mice, 8. Bars show geometric mean ± 95% C.I. Dots indicate repeats.

(B-D, E, G) Bars show mean ± s.e.m. Dots indicate repeats. *, $p < 0.05$; **, $p < 0.01$; ***, $p < 0.001$ by Dunett's test with CTRL or T-test.

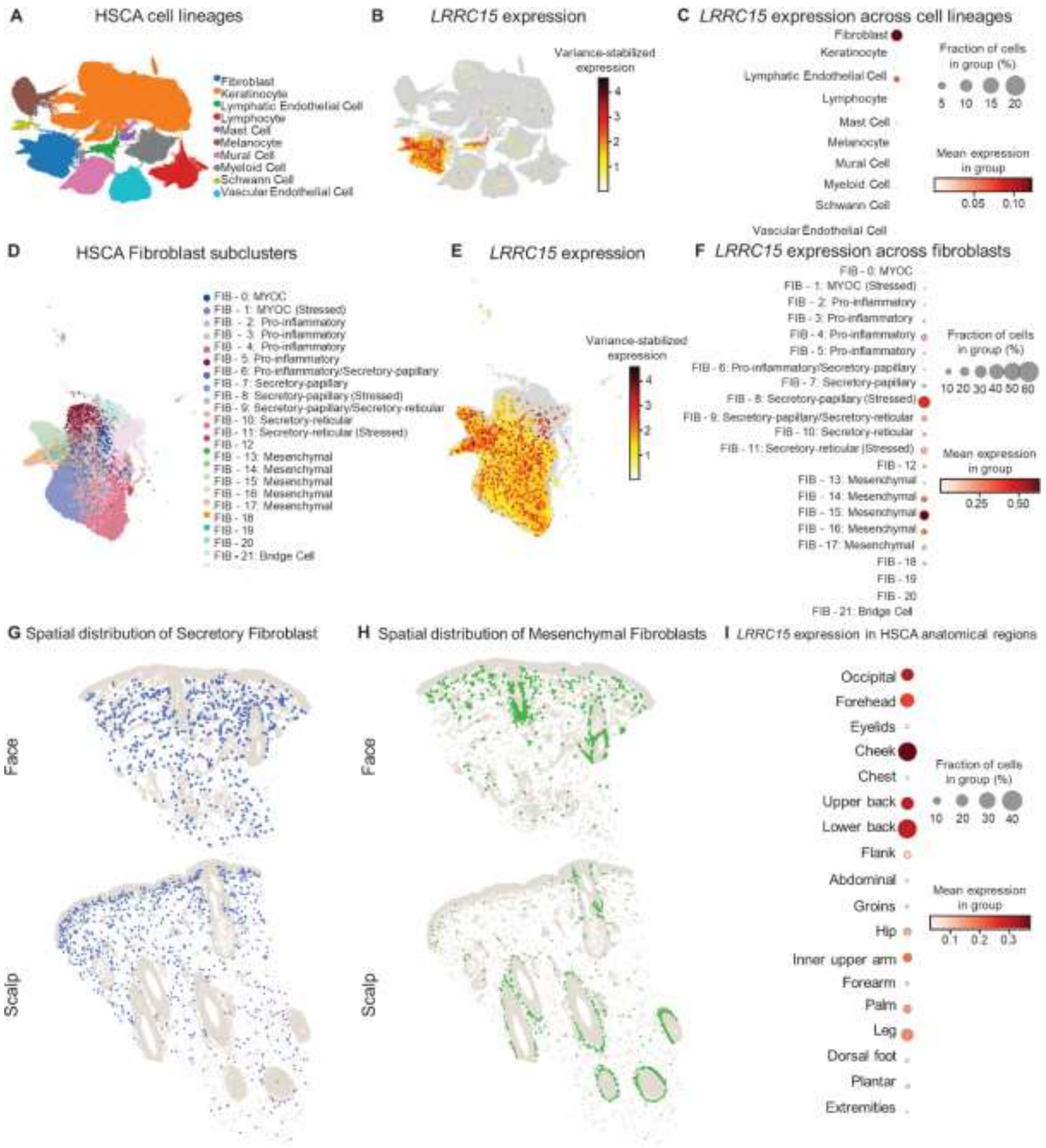

**Figure S11. Meta-analysis of *LRRC15* expression in skin Human Cell Atlas (HCA), related to Figure 6**

**(A)** UMAP plot of present cell lineages in HCA.

**(B)** Feature expression plot of *LRRC15* across all cells.

**(C)** Dot plot of average *LRRC15* expression across cell lineages.

**(D)** UMAP of fibroblast subclusters.

**(E)** Feature expression plot of *LRRC15* within fibroblast subclusters.

**(F)** Dot plot of average *LRRC15* expression across all fibroblast subclusters.

**(G–H)** Spatial mapping of HCA secretory fibroblasts (G) and HCA mesenchymal fibroblasts (H) onto Xenium spatial transcriptomics data generated from face and scalp samples.

**(I)** Dot plot of average *LRRC15* expression in fibroblasts across sampled anatomical sites in HCA.

**Table S1. Primer list**

| **Target gene** | **Primer ( 5' → 3')** | **Primer (5' → 3')** | **References** |
|---|---|---|---|
| Envelop E of WNV-IS98 | ATTCGGGAGGAGACGTGGTA | CAGCCGCCAACATCAACAAA | This study |
| *Saa2* | TACAGCACAGATCAGCACCA | CTCAGGACCAAGGAGCAGAA | |
| *Defb1* | TGTCTCTATTCTGCCTGCCC | CACTTGGCCTTCCCTCTGTA | |
| *Lrrc15* | CTCAATGAGTCCCCGTTCCT | TTGTTGGCGAGGCTGAGATA | |
| *Gapdh* | AAATCAAGTGGGGCGATGCT | AAGCAGTTGGTGGTGCAGGA | Deramaudt et al., 2006 [3] |
| *Actb* | AGCACAGAGCCTCGCCTT | CATCATCCATGGTGAGCTGG | Maarifi G *et al.*, 2021 [4] |
| *Rpl13a* | AACAGCTCATGAGGCTACGG | TGGGTCTTGAGGACCTCTGT | |
| *Lrrc15* (Mouse) | AACACACACATCACCGAACTC | AGATTGCGGAAGGCACCTGG | Toriumi K *et al.*, 2023 [5] |

**Dataset S1. Clinical scores for mouse experiments, related to Figures 1 and 6**

**A**. Clinical score interpretation.

**B**. Clinical score for Figure S1D.

**C**. Clinical score for Figure 5Q.

**Dataset S2. Cell type-enriched genes, related to Figure 1**

**Dataset S3. Cell subtype-enriched genes, related to Figure 1**

**Dataset S4. Differentially expressed genes in mosquito RNA-containing fibroblasts, related to Figure 2**

**Dataset S5. Differentially expressed genes in infectious bite and their functional classification, related to Figure 5**
